# The dependence of EGFR oligomerization on environment and structure: A camera-based N&B study

**DOI:** 10.1101/2022.05.06.490852

**Authors:** Harikrushnan Balasubramanian, Jagadish Sankaran, Corinna Jie Hui Goh, Thorsten Wohland

**Affiliations:** Department of Biological Sciences and NUS Centre for Bio-Imaging Sciences, National University of Singapore, Singapore; Department of Chemistry, National University of Singapore, Singapore

## Abstract

Number and Brightness analysis (N&B) is a fluorescence spectroscopy technique to quantify protein oligomerization. Accurate results, however, rely on a good knowledge of non-fluorescent states of the fluorescent labels, especially of fluorescent proteins (FP), which are widely used in biology. FPs have been characterized for confocal but not camera-based N&B, which allows in principle faster measurements over larger areas. Here, we calibrate camera-based N&B implemented on a total internal reflection fluorescence (TIRF) microscope for various fluorescent proteins by determining their propensity to be fluorescent. We then apply camera-based N&B in live CHO-K1 cells to determine the oligomerization state of the epidermal growth factor receptor (EGFR), a transmembrane receptor tyrosine kinase that is a crucial regulator of cell proliferation and survival with implications in many cancers. EGFR oligomerization in resting cells and its regulation by the plasma membrane microenvironment is still under debate. Therefore, we investigate the effects of extrinsic factors, including membrane organization, cytoskeletal structure, and ligand stimulation, and intrinsic factors, including mutations in various EGFR domains, on the receptor’s oligomerization. Our results demonstrate that EGFR oligomerization increases with removal of cholesterol or sphingolipids, or the disruption of GM3-EGFR interactions, indicating raft association. However, oligomerization was not significantly influenced by the cytoskeleton. Mutations in either I706/V948 residues or E685/E687/E690 residues in the kinase and juxtamembrane domains, respectively, led to a decrease in oligomerization, indicating their necessity for EGFR dimerization. Finally, EGFR phosphorylation is oligomerization-dependent involving the extracellular domain (550-580 residues). Coupled with biochemical investigations, camera-based N&B indicates that EGFR oligomerization and phosphorylation is the outcome of several molecular interactions involving the lipid content and structure of the cell membrane and multiple residues in the kinase, juxtamembrane, and extracellular domains.

**STATEMENT OF SIGNIFICANCE:** Number and brightness (N&B) analysis is a powerful tool to determine protein association but is mostly conducted in confocal microscopes. This work determines the brightness and fluorescence probability of a range of fluorescent proteins for camera-based N&B on a total internal reflection microscope, demonstrating that with proper calibration different fluorescent proteins provide the same answers on oligomerization within the margins of error. This camera-based approach allows measuring N&B values across whole cell basal membranes up to an area of ~1,000 μm^2^ simultaneously. N&B is then used in combination with biochemical assays to investigate the oligomerization and activation of the epidermal growth factor receptor (EGFR), a prototypical receptor tyrosine kinase with importance in cell signalling, division and survival and implicated in various cancers. The results indicate that EGFR oligomerization and activation is governed by an interplay between membrane structure and composition and key amino acid residues of EGFR that span the extracellular to the intracellular domains.

## INTRODUCTION

Protein oligomerization or clustering is an essential process in many cell signalling pathways and its determination and quantification in living cells is an important task in microscopy. More often than not, biological systems contain not a single species but a mixture of various oligomeric species that contribute differently to the signalling process and their separation is difficult. Multiple techniques based on fluorescence spectroscopy including Förster resonance energy transfer (FRET) (1, 2), fluorescence cross-correlation spectroscopy (FCCS) (3), photon counting histogram (PCH) (4), fluorescence intensity distribution analysis (FIDA) (5), spatial intensity distribution analysis (SpIDA) (6, 7), Image Correlation Spectroscopy (ICS) (8), and single-molecule photobleaching (SMP) (9) have been developed to address this challenge, each having their own set of requirements, advantages, and disadvantages.

One of the techniques to quantify oligomerization is number and brightness analysis (N&B) (10, 11), a fluorescence fluctuation spectroscopy method based on moment analysis (12, 13). In this technique, the average number and brightness of particles is calculated from the mean and variance, i.e., the first and second moments, respectively, of the intensity fluctuations. N&B was originally established for confocal microscopy (10, 11) and later expanded to camera-based total internal reflection microscopy (TIRFM) (14). Due to its computational simplicity, it is widely used for oligomerization studies of various biological molecules, including cell membrane receptors (15).

N&B is typically performed on proteins of interest fused with fluorescent proteins (FP). The oligomerization state of a protein is determined by computing the ratio of the brightness of the protein of interest to the brightness of the monomeric state of the FP. However not all molecules of FPs are fluorescent due to incomplete maturation, misfolding or photobleaching (16–20). Therefore, N&B is calibrated with singly and doubly labelled biomolecules. The singly labelled sample provides a direct measure of the molecular brightness of the label. In principle the doubly labelled sample should have twice the brightness. Deviation from this yields the probability of the label to be fluorescent. Here, we first determine the brightness and fluorescence probability of cell membrane-bound FPs from the fluorescent protein toolbox (21) and apply it to the investigation of the oligomerization of the epidermal growth factor receptor (EGFR).

EGFR is a receptor tyrosine kinase that transduces signals through phosphorylation of intracellular tyrosine residues. Although EGFR has been one of the most-widely studied proteins, its resting state conformation remains a matter of debate. The traditional view is that EGFR exists as monomers on the cell membrane prior to ligand binding and undergoes dimerization upon EGF binding. However, pre-formed EGFR dimers in the absence of EGF have been reported in numerous studies (Figure S1) (5, 22–28). But the dimerization levels vary greatly between studies (Table S1), and it is unclear what are the determining factors for receptor di- or oligomerization (26, 29). Furthermore, there is evidence that full receptor activation requires higher oligomers (28, 30–35). Here we use camera-based N&B on TIRFM to determine the oligomerization of EGFR in live CHO-K1 cells and investigate the effects of the actin cytoskeleton, the plasma membrane organization, and the roles of different EGFR residues on oligomerization.

## THEORY

### N&B

The N&B equations for apparent number and apparent brightness using an EMCCD are (14)

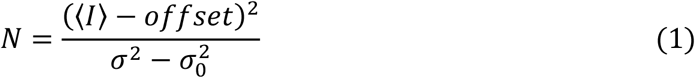

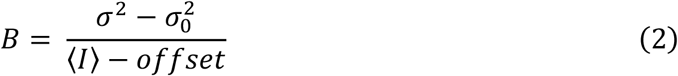

where *N* is the apparent number of particles in the observation volume, *B* is the apparent brightness of a particle, 〈*I*〉 is the average signal intensity, *offset* is the intensity offset of the camera, *σ*^2^ is the variance of the signal, and 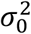 is the variance of the readout noise in the camera. The variance of the signal is strongly influenced by shot noise, and thus we estimate the true variance from the covariance between consecutive frames, an approach called G1 analysis (Eq. S1) (14). This is analogous to the estimation of the amplitude of the autocorrelation function, G(0), in fluorescence correlation spectroscopy, from neighbouring values.

### Workflow of N&B

The schematic in Figure 1 shows the workflow of the TIRF-implemented camera-based N&B used in this study. It illustrates with an example the power of N&B to identify the proportional contribution of the *N* and *B* values to the pixel intensity from its mean and variance alone (Eqs. 1 and 2).

**Figure 1:**
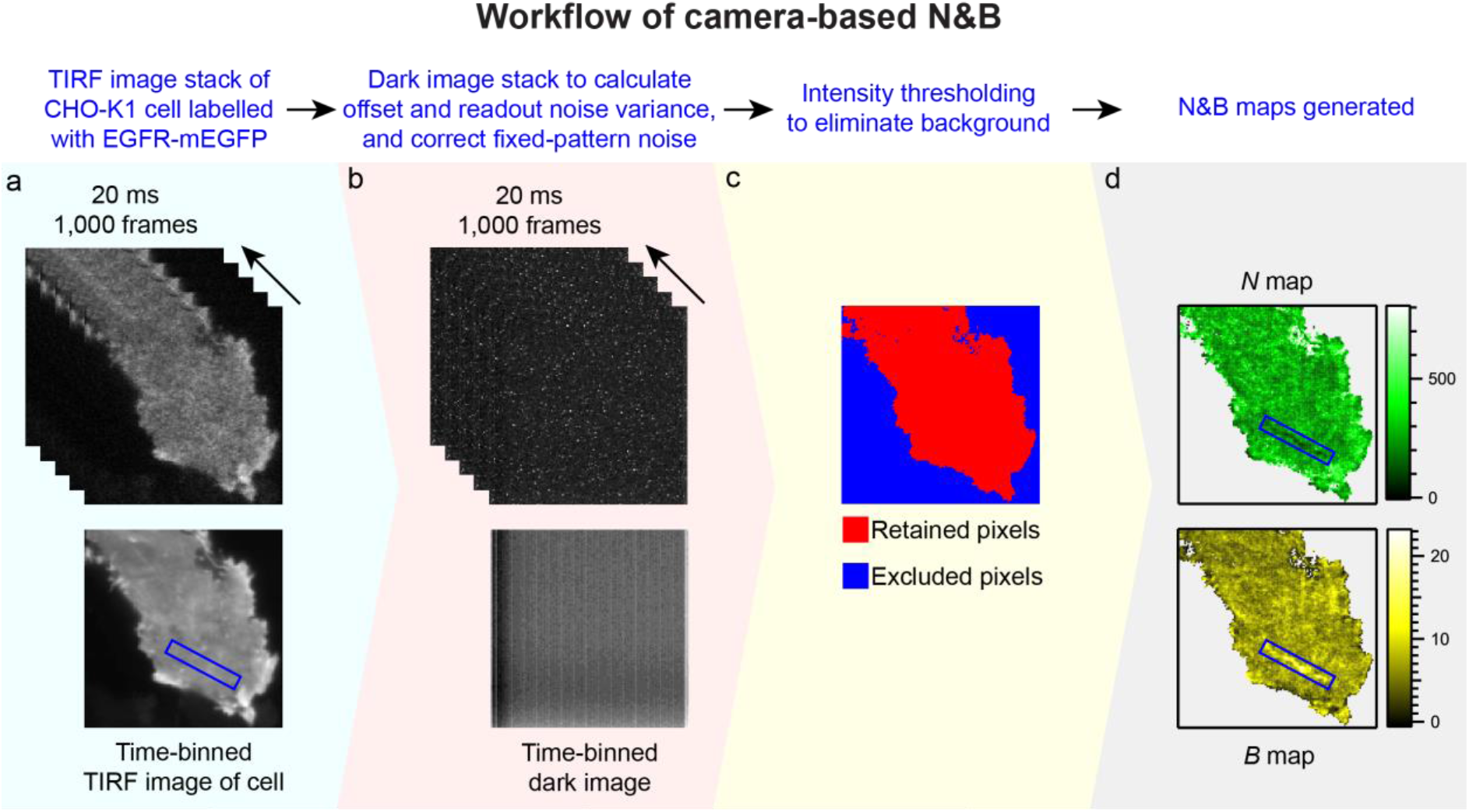
N&B workflow. **(a)** A CHO-K1 cell labelled with EGFR-mEGFP is shown as an example. The TIRF image stack of this cell contains 1,000 frames collected at 20 ms exposure time. The TIRF image after time-binning of all the frames is shown at the bottom left. **(b)** A dark image stack from the camera with the same spatiotemporal characteristics as the cell image stack is used to obtain the offset and readout noise variance and to correct for fixed-pattern noise, as shown by the projection of all frames in the bottom image. **(c)** An intensity threshold (refer to *Data analysis* section in *Materials and Methods*) is applied to exclude the background pixels (blue) and retain only pixels from the cell (red). **(d)** Applying Eqs. 1 and 2, *N* and *B* maps are generated. Although, the blue box in the time-binned TIRF image shows an area whose intensity is indistinguishable from the surrounding areas, the *N* and *B* maps of the same area show that the number of particles is low and the molecular brightness is high, respectively, illustrating the utility of N&B.

The *offset* and 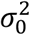 is obtained from a dark image captured by the camera. The dark image also serves as a reference image to perform pixel-wise correction of fixed-pattern noise of camera images (supplement section 1.2).

### Pooled brightness of cells

For a population of cells, the pooled weighted arithmetic mean brightness is calculated from the brightness values of pixels in cell images (27). Each image stack contains only one cell.

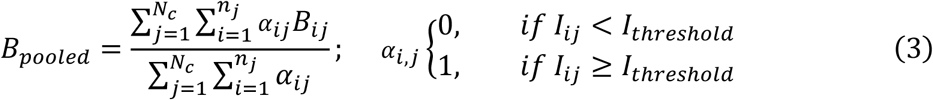

where *B_pooled_* is the pooled mean brightness for a population of *N_c_* cells, *n_j_* is the number of pixels in image *j*, and *B_ij_* is the brightness of each pixel *i* in image *j. α_ij_* is 1 if the intensity of a pixel (*I_ij_*) is more than the intensity threshold value (*I_threshold_*) and corresponds to pixels inside a cell and vice-versa. The error in *B_pooled_* is calculated according to Eq. S3. In the rest of the manuscript all brightness values are pooled values.

### Brightness calibration

The theory of N&B for oligomers is published in detail in a previous publication (27). Here and in the supporting material we provide a brief summary. The fluorescence probability *p* of a fluorescent protein is estimated from the ratio of the pooled apparent brightness of the dimeric FP control (*B_d_*) to the pooled apparent brightness of the monomeric FP control (*B_m_*) (27).

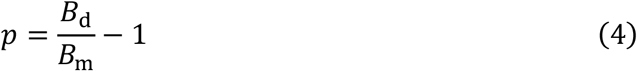

The error in *p* is calculated according to Eq. S4.

### EGFR oligomerization calculations

In the case of EGFR experiments, we first calculate the ratio of the apparent brightness of EGFR (*B*_E_) to the apparent brightness of the monomer (*B*_m_).

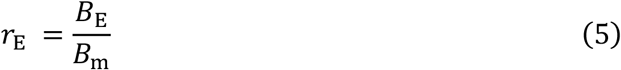

We utilize the minimum-order oligomer model (MOM) to determine whether EGFR is monomeric or whether oligomers are necessary to explain the data (27). The MOM model assumes the presence of only two species – monomers and the minimum-order oligomer species required to explain the data. This leaves open the question whether a mixture of more and higher oligomer species are present. But it provides a lower limit of oligomerization state and an upper limit of the fraction of monomers bound in oligomers. Using MOM, the fraction of EGFR molecules present as oligomers (*e_oligo_*, refer to Figure S3) is expressed as (27):

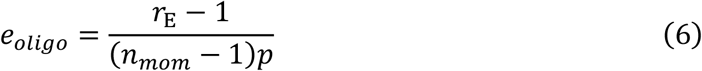

where *n_mom_* is the order of the minimum oligomer species. In order to determine *n_mom_*, we estimate the brightness of pure higher order oligomer samples (*B_n_*) based on the experimental *B* and *p* values of the monomer and dimer FP controls (Figure S4) (27).

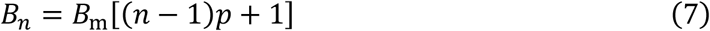

where *n* is the oligomer species order. The errors in *r*_E_, *e_oligo_* and *B_n_* are calculated according to Eqs. S5, S6 and S7, respectively.

The minimum value of *n* yielding *B_n_* > *B_E_* is used as *n_mom_* where ⌈ ⌉ is the ceiling function.

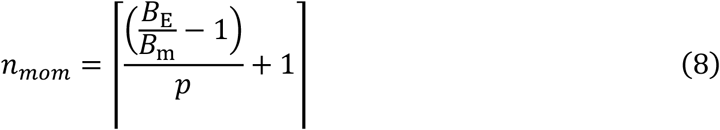

Once the minimum order is determined, the proportion of the monomers and the minimum order oligomers is determined using Eq. 6 (27). While MOM may not represent the true distribution of various oligomer species, it provides apparent oligomer percentages with the understanding that they represent a minimal model of monomers and the smallest oligomer species that are consistent with the experimental data.

We define two terms, oligomerization and clustering, as follows. Oligomerization involves specific receptor-receptor interactions to form larger oligomeric units which activate the receptor. Oligomers are typically below the diffraction limit and cannot be optically resolved. Even in areas of homogenous intensity in images, certain pixels of *B* values larger than the dimer calibration control are present which are indicative of oligomers. Clustering may involve oligomerization of EGFR, grouping of receptors localized in lipid domains, non-specific aggregation of receptors, and most likely a combination of all three. Clusters are seen as large and clearly-defined visible patches of multiple pixels in images (e.g., refer blue box in Figure 1).

## MATERIALS AND METHODS

### Sample preparation

Details of plasmids, preparation of live cell samples, drug perturbations, and western blotting are provided in the Supporting Material. Full protocols for live cell sample preparation and data acquisition and analysis are also available on Protocol Exchange (36, 37).

### Instrumentation

The TIRF microscopy set-up included an inverted epifluorescence microscope (IX83, Olympus, Japan) equipped with a motorized TIRF illumination combiner (cellTIRF-4Line IX3-MITICO, Olympus, Japan). 405 nm (LAS/405/100/D, Olympus, Japan), 488 nm (LAS/488/100, Olympus, Japan) and 561 nm (LAS/561/100, Olympus, Japan) lasers were connected to the TIRF illumination combiner. The 488 nm laser was equipped with a clean-up filter (FF01-488/6-25, Semrock, New York, USA). A 100×, NA 1.49 oil-immersion objective (Apo N, Olympus, Japan) was used. The immersion oil used had a refractive index of 1.518 (IMMOIL-F30CC, Olympus, Japan). A quad band dichroic (ZT405/488/561/640rpc, Chroma Technology Corp, Vermont, USA) was used to direct the laser to the sample and allow the fluorescence emission to pass through. The fluorescence emission was passed through a quad band emission filter (ZET405/488/561/640m, Chroma Technology Corp, Vermont, USA). An electron-multiplying charge-coupled device (EMCCD; iXon^EM^+ 860, 24 μm pixel size, 128 × 128 pixels, Andor, Oxford Instruments, UK) camera was used for detection. For the cell measurements, 37°C temperature and 5% CO_2_ atmosphere were maintained using an on-stage incubator (Chamlide TC, Live Cell Instrument, South Korea).

### Data acquisition

We used 100 μW power (as measured in the back focal plane of the objective) for all lasers. The penetration depth was set at 100 nm. The software Andor Solis (version 4.31.30037.0-64-bit) was used for image acquisition. The following camera settings were used for the EMCCD: mode of image acquisition = kinetic, baseline clamp (i.e., offset) = “on” to minimize the baseline fluctuation, pixel readout speed = 10 MHz, maximum analog-to-digital gain = 4.7, vertical shift speed = 0.45 μs, electron multiplying (EM) gain = 300. The EMCCD was operated after cooling to −80°C. A stack of 1,000 frames (128 × 128 pixels) were collected at 50 frames per second. The recorded image stacks were saved as 16-bit tiff files. A dark image was collected with the camera shutter closed using the same acquisition parameters.

### Data analysis

The data analyses were performed on a computer equipped with a GPU with the following configuration: Windows 10 Home 64-bit operating system, Intel® Core™ i7-7800X CPU @ 3.50 GHz processor, 32 GB RAM, NVIDIA TITAN Xp GPU with 3,840 CUDA cores and 12.3 GB memory. The image stacks were loaded into the GPU-driven ImFCS 1.52 plugin (27, 38) in Fiji. A polynomial of order 2 was used to correct for photobleaching. Intensity thresholding was performed to include only pixels within the cell. Background correction was performed using a dark image with the same spatiotemporal dimensions as the measurement image.

## RESULTS AND DISCUSSION

### Brightness calibration for N&B

#### Optimization of acquisition parameters for N&B

N&B requires an exposure time that is shorter than the time a particle needs to diffuse through the observation area of a pixel but long enough to obtain a stable N&B signal with high SNR (27, 39). This time was determined to be 20 ms for our camera-based system (27). Moreover, an exposure time of 20 ms allowed better intensity thresholding of the pixels to separate the cell from the background.

The PMT-mEGFP_2_/PMT-mEGFP brightness ratio measured at 20 ms exposure time was determined for total acquisition times from 1-100 s. It is variable below 10 s and then stabilised at ~1.8 (Figure S5). In this work, we used 20 s for all measurements as a compromise between acquisition speed, and accuracy and precision. Measurements had photobleaching, i.e., a steady decay of intensity, that influences that variance and thus the calculated *N* and *B* values. Therefore, a photobleaching correction, using a polynomial of 2^nd^ degree (40, 41), is applied to avoid influence of the decaying intensity on N&B, very similar to fluorescence correlation spectroscopy (15, 42, 43). Example *B* maps and intensity traces for PMT-mEGFP, PMT-mEGFP_2_, and EGFR-mEGFP are shown in Figure S6 to illustrate the effect of photobleaching on *B* values.

#### Brightness calibration of various fluorescent proteins

We performed brightness calibration for seven different FPs (mTurquoise2 (51), mEGFP (52), mApple (53), mCherry (54), mCherry2 (55), mScarlet (48), and mKate2 (56)). It is critical that the FPs used are predominantly monomeric and do not have a tendency to form homo-oligomers. The monomericities as reported in literature for the various FPs tested in this study are provided in Table 1. mEGFP exhibits a fluorescence probability *p* of 0.81 ± 0.01, which is the highest among all the tested FPs. Among the other FPs, *p* varied between 30-65%. These *p* values are consistent with those reported in previous studies using confocal microscopy (16, 18).

**Table 1:** Brightness calibration of various fluorescent proteins.

| Fluorescent protein | Monomericity reported in literature | Brightness calibration control | Cells | $B^a$ | $p^a$ |
| --- | --- | --- | --- | --- | --- |
| mTurquoise2 | 88-94%<br>(17, 44) | PMT-mTurquoise2 | 17 | $1.83 \pm 0.02$ | $0.39 \pm 0.02$ |
| | | PMT-mTurquoise2<sub>2</sub> | 18 | $2.54 \pm 0.02$ | |
| mEGFP | 97-98%<br>(17, 45–47) | PMT-mEGFP | 21 | $2.32 \pm 0.01$ | $0.81 \pm 0.01$ |
| | | PMT-mEGFP<sub>2</sub> | 18 | $4.21 \pm 0.02$ | |
| mApple | 87-95%<br>(17, 48) | PMT-mApple | 20 | $1.38 \pm 0.03$ | $0.53 \pm 0.03$ |
| | | PMT-mApple<sub>2</sub> | 21 | $2.11 \pm 0.02$ | |
| mScarlet | 80%<br>(48) | PMT-mScarlet | 19 | $3.74 \pm 0.03$ | $0.54 \pm 0.03$ |
| | | PMT-mScarlet<sub>2</sub> | 15 | $5.75 \pm 0.02$ | |
| mCherry | 69-95%<br>(17, 48–50) | PMT-mCherry | 16 | $0.86 \pm 0.04$ | $0.32 \pm 0.03$ |
| | | PMT-mCherry<sub>2</sub> | 18 | $1.14 \pm 0.04$ | |
| mCherry2 | Not reported in literature yet | PMT-mCherry2 | 15 | $0.83 \pm 0.04$ | $0.64 \pm 0.03$ |
| | | PMT-mCherry2<sub>2</sub> | 16 | $1.36 \pm 0.04$ | |
| mKate2 | 59-81%<br>(17, 48) | PMT-mKate2 | 17 | $1.17 \pm 0.04$ | $0.52 \pm 0.04$ |
| | | PMT-mKate2<sub>2</sub> | 18 | $1.78 \pm 0.04$ | |

<sup>a</sup> The reported values for $B$ and $p$ are in terms of mean ± SEM.
The number of pixels in each cell is different due to differential intensity filtering in each case, but each cell had at least 3,000 and at most 17,000 valid pixels.

In theory, knowledge of the *p* value allows any FP of choice to be used for obtaining reliable oligomerization results, as we demonstrated comparing mEGFP and mApple (27). But as can be seen from Table 1 many FPs considered monomeric are not truly so. FP homo-oligomerization affects the brightness calibration and also introduces uncertainty in the oligomerization estimates of the protein under investigation. Moreover, FPs with higher *p* values are better for visualization of labelled samples since more molecules are fluorescent. In this study we used mEGFP which is highly monomeric (97-98% (17, 47)) and has high fluorescence probability (*p* = 0.81 ± 0.01).

### EGFR oligomerization in resting state determined by N&B

In resting cells, the mean brightness (*B*_E_) of EGFR-mEGFP is 3.39 ± 0.02 (Table 2). The *B* values of the calibration controls PMT-mEGFP and PMT-mEGFP_2_ are 2.32 ± 0.01 and 4.21 ± 0.02 (Table 1). Since 81% of mEGFP molecules are fluorescent, *r*_E_ (refer to Eq. 5) < 1.81 can be explained by a dimerization model while a higher oligomerization model is required to explain *r*_E_ > 1.81. Here, *r*_E_ = 1.46 is explained using a dimerization model. This *r*_E_ value suggests the dimerization of 57 ± 2% of EGFR molecules in resting cells. This is in agreement with 50-70% dimerization reported in the resting state for the same cell line in previous studies (5, 25, 26, 28, 57). The ~60% dimerization observed is similar to what we had observed in our previous publication (27) where we tagged EGFR with mApple.

**Table 2:**
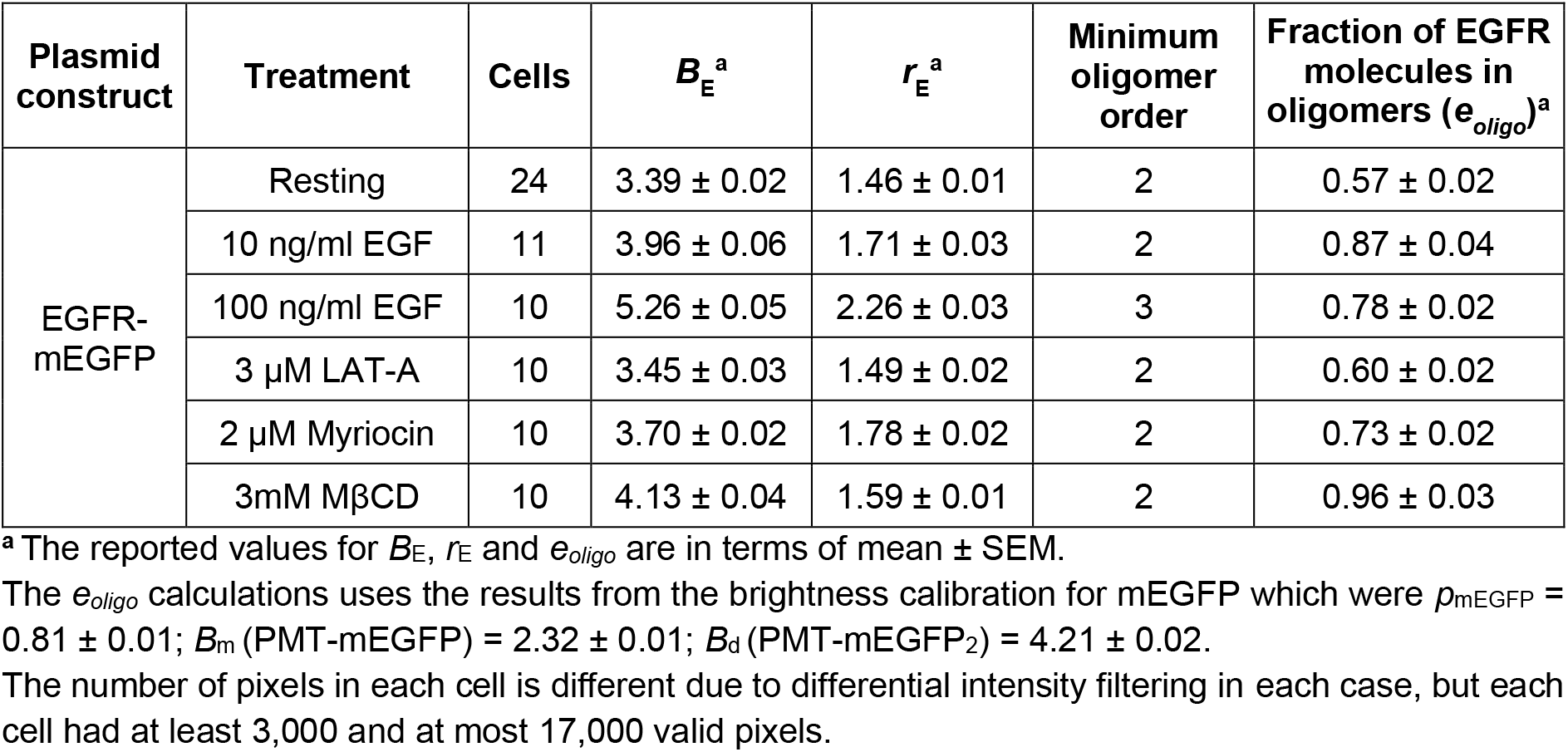
N&B of EGFR-mEGFP in resting state, after ligand stimulation and when subjected to various drug treatments.

| Plasmid construct | Treatment | Cells | $B_E^a$ | $r_E^a$ | Minimum oligomer order | Fraction of EGFR molecules in oligomers ($e_{oligo}$)<sup>a</sup> |
| --- | --- | --- | --- | --- | --- | --- |
| EGFR-mEGFP | Resting | 24 | $3.39 \pm 0.02$ | $1.46 \pm 0.01$ | 2 | $0.57 \pm 0.02$ |
| | 10 ng/ml EGF | 11 | $3.96 \pm 0.06$ | $1.71 \pm 0.03$ | 2 | $0.87 \pm 0.04$ |
| | 100 ng/ml EGF | 10 | $5.26 \pm 0.05$ | $2.26 \pm 0.03$ | 3 | $0.78 \pm 0.02$ |
| | 3 μM LAT-A | 10 | $3.45 \pm 0.03$ | $1.49 \pm 0.02$ | 2 | $0.60 \pm 0.02$ |
| | 2 μM Myriocin | 10 | $3.70 \pm 0.02$ | $1.78 \pm 0.02$ | 2 | $0.73 \pm 0.02$ |
| | 3mM MβCD | 10 | $4.13 \pm 0.04$ | $1.59 \pm 0.01$ | 2 | $0.96 \pm 0.03$ |

<sup>a</sup> The reported values for $B_E$ , $r_E$ and $e_{oligo}$ are in terms of mean ± SEM.
The $e_{oligo}$ calculations uses the results from the brightness calibration for mEGFP which were $p_{mEGFP} = 0.81 \pm 0.01$ ; $B_m$ (PMT-mEGFP) = $2.32 \pm 0.01$ ; $B_d$ (PMT-mEGFP<sub>2</sub>) = $4.21 \pm 0.02$ .
The number of pixels in each cell is different due to differential intensity filtering in each case, but each cell had at least 3,000 and at most 17,000 valid pixels.

Intensity counts of the images are a qualitative proxy for expression levels. Over a range of intensities from 500 to 8000 counts, the average *B* values of the cells expressing EGFR-mEGFP (*n*=10, Figure S7) do not vary with EGFR expression levels. This implies that these expression levels do not influence EGFR clustering in our experiments. This is similar to previous studies (26, 57) which also found no dependence of dimerization on the expression levels of EGFR in cells expressing in the range of 10,000 to 1.6 million receptors per cell. It is to be noted that since cameras are used in this study and they are not photon-counting devices, absolute expression levels are not known (42).

### EGFR oligomerization after low- and high-dose EGF stimulation

Low- and high-dose ligand stimulation of EGFR was achieved using 10 ng/ml and 100 ng/ml EGF, respectively (Figure 2). In both cases, there was an increase in *B*_E_ by 1.17-fold and 1.55-fold, respectively, from the resting state. The *B*_E_ values of 3.96 ± 0.06 (10 ng/ml EGF) and 5.26 ± 0.05 (100 ng/ml EGF) (Table 2) correspond to *r*_E_ values of 1.71 and 2.27, respectively. In the case of 10 ng/ml EGF stimulation, *r*_E_ = 1.71 is explained by a dimerization model and translates to 87 ± 4% of EGFR receptors forming dimers.

**Figure 2:**
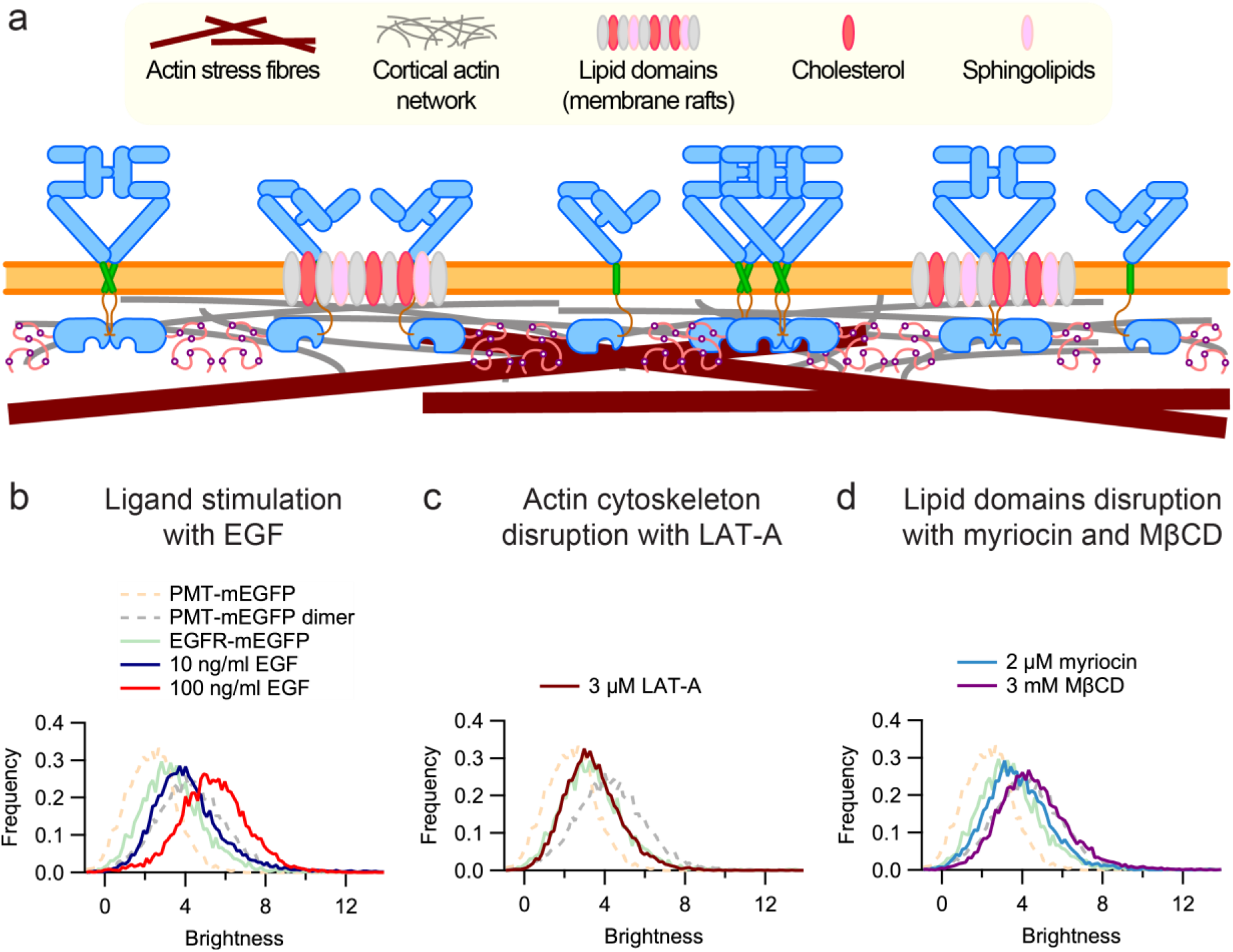
Brightness distributions of EGFR-mEGFP after ligand stimulation and drug perturbations. **(a)** This schematic shows EGFR distribution on the plasma membrane. The underlying actin cytoskeleton (stress fibres and cortical actin) provides stability to the plasma membrane. EGFR is present as monomers, dimers and oligomers. A fraction of the EGFR population is trapped in lipid domains. Representative brightness histograms are shown here for the calibration controls (light pink and grey dotted curves), EGFR-mEGFP in resting state (green curve), and the results of the drug treatments – **(b)** EGF stimulation (blue and red curves), **(c)** actin cytoskeleton disruption with LAT-A (brown curve), and **(d)** disruption of lipid domains with myriocin (blue curve) and MβCD (purple curve).

In the case of 100 ng/ml EGF stimulation, an *r*_E_ = 2.27 cannot be explained by a dimerization model. An oligomerization model is invoked to explain the *r*_E_ value. Using MOM, the minimum order oligomer required to explain *r*_E_ = 2.27 is a trimer. Hence, this *r*_E_ value translates to 78 ± 2% of EGFR molecules forming trimers. Note that this does not imply that only EGFR monomers and trimers are present, or that trimers are the dominant oligomeric species. It is very likely that several oligomeric species are present (5, 28) and some studies using homo-FRET have suggested that tetramers are the dominant oligomeric species (30–32, 58).

### Effects of plasma membrane components on EGFR oligomerization

EGFR oligomerization was investigated after perturbation of the cells with various drugs (refer to Supplemental materials and methods) that are known to affect EGFR organization and functioning. The results from all these measurements are shown in Table 2 and representative schematics and brightness histograms are shown in Figure 2.

#### EGFR oligomerization after actin depolymerization using LAT-A

After treatment with 3 μM latrunculin A (LAT-A) to test the effects of actin depolymerization on EGFR clustering (Figure 2), we measured a *B*_E_ of 3.45 ± 0.03 (Table 2). The value of *r*_E_ of 1.49 thus remained similar to the resting state with an *r*_E_ of 1.46, indicating that the cytoskeleton does not influence EGFR oligomerization. Our results are consistent with studies using FCCS and fluorescence lifetime imaging-FRET (FLIM-FRET) (26, 59).

#### EGFR oligomerization after cholesterol depletion using MβCD

Numerous studies have reported that a part of the EGFR population is trapped in lipid domains (5, 26, 60–66). Since lipid domains are enriched in cholesterol (67), cholesterol depletion disrupts these domains. We treated cells with 3 mM methyl-β-cyclodextrin (MβCD) to investigate the effect of cholesterol depletion on EGFR oligomerization (Figure 2). After treatment the *B*_E_ value increased to 4.13 ± 0.04 (Table 2) which is a 1.2-fold increase from the resting state. The *r*_E_ value of 1.78 indicates at least strong dimerization with 96 ± 3% of EGFR molecules dimerizing.

These results suggest that EGFR oligomerization is hindered by trapping of EGFR molecules in lipid domains. Depletion of cholesterol perturbs these domains and releases the trapped EGFR molecules. The freed receptors interact with each other to form oligomers. Other studies have also reported an increase in EGFR oligomerization and phosphorylation after cholesterol depletion using both biochemical (cross-linking, immunoprecipitation) and biophysical (FCCS, SMP, immunoelectron microscopy) techniques (5, 26, 60–66).

#### EGFR oligomerization after sphingolipids depletion using myriocin

Sphingolipids are also a core component of lipid domains (67) and represent an alternative target to perturb these domains. The importance of sphingolipids in EGFR clustering was investigated using 2 μM myriocin which interferes with sphingolipid synthesis (Figure 2) (67, 68). We measured a *B*_E_ value of 3.70 ± 0.02 (Table 2) which is a 1.09-fold increase compared to the resting state with a concomitant value of *r*_E_ = 1.60. This data can be explained by 73 ± 2% of EGFR molecules existing in dimers. This indicates that myriocin treatment is able to free a fraction of the receptors from the lipid domains and increases their probability to interact and oligomerize, although its effect is not as strong as that of MβCD (69).

### Effects of EGFR residues on EGFR oligomerization

Here we investigate the importance of various regions in EGFR molecule in EGFR clustering by creating point and deletion mutants in those regions. The mutants were tested in the resting state and after low- and high-dose EGF stimulation.

Four mutants were created: EGFR^I706Q/V948R^-mEGFP (70), EGFR^E685A/E687A/E690A^-mEGFP (71), EGFR^K642G^-mEGFP (72), and EGFR^Δ550-580^-mEGFP (73). The results from the N&B of these mutants are shown in Table 3 and representative brightness histograms are shown in Figure 3. Furthermore, western blot-based phosphorylation assays were conducted to corroborate the N&B data from a biochemical perspective and the results are shown in Figure S8.

**Table 3:** N&B of EGFR-mEGFP and EGFR-mEGFP mutants in resting state and after EGF stimulation.

| Mutation | Treatment | Cells | $B_E^a$ | $r_E^a$ | Minimum oligomer order | Fraction of EGFR molecules in oligomers ($e_{oligo}$)<sup>a</sup> | Relative phosphorylation (Figure S8) |
| --- | --- | --- | --- | --- | --- | --- | --- |
| EGFR-mEGFP | Resting | 24 | 3.39 ± 0.02 | 1.46 ± 0.01 | 2 | 0.57 ± 0.02 | 0 |
|  | 10 ng/ml EGF | 11 | 3.96 ± 0.06 | 1.71 ± 0.03 | 2 | 0.87 ± 0.04 | NA |
|  | 100 ng/ml EGF | 10 | 5.26 ± 0.05 | 2.26 ± 0.03 | 3 | 0.78 ± 0.02 | 1 |
| I706Q/V948R | Resting | 28 | 2.88 ± 0.02 | 1.24 ± 0.01 | 2 | 0.29 ± 0.01 | 0.1 |
|  | 10 ng/ml EGF | 10 | 3.20 ± 0.02 | 1.38 ± 0.01 | 2 | 0.46 ± 0.02 | NA |
|  | 100 ng/ml EGF | 10 | 3.36 ± 0.02 | 1.44 ± 0.01 | 2 | 0.55 ± 0.02 | 0.1 |
| E685A/E687A/E690A | Resting | 28 | 2.69 ± 0.02 | 1.46 ± 0.01 | 2 | 0.19 ± 0.01 | 0 |
|  | 10 ng/ml EGF | 12 | 3.33 ± 0.03 | 1.16 ± 0.01 | 2 | 0.53 ± 0.02 | NA |
|  | 100 ng/ml EGF | 13 | 4.69 ± 0.04 | 1.43 ± 0.02 | 3 | 0.63 ± 0.02 | 1.8 |
| K642G | Resting | 21 | 4.11 ± 0.03 | 1.77 ± 0.02 | 2 | 0.95 ± 0.03 | 0.4 |
|  | 10 ng/ml EGF | 13 | 4.73 ± 0.03 | 2.04 ± 0.02 | 3 | 0.64 ± 0.02 | NA |
|  | 100 ng/ml EGF | 12 | 5.53 ± 0.06 | 2.38 ± 0.03 | 3 | 0.85 ± 0.02 | 6.4 |
| Δ550-580 | Resting | 21 | 3.18 ± 0.02 | 1.37 ± 0.01 | 2 | 0.45 ± 0.02 | 0.2 |
|  | 10 ng/ml EGF | 10 | 3.40 ± 0.03 | 1.47 ± 0.02 | 2 | 0.58 ± 0.01 | NA |
|  | 100 ng/ml EGF | 10 | 3.64 ± 0.02 | 1.57 ± 0.01 | 2 | 0.70 ± 0.02 | 0.1 |

<sup>a</sup> The reported values for $B_E$ , $r_E$ and $e_{oligo}$ are in terms of mean ± SEM.
The $e_{oligo}$ calculations uses the results from the brightness calibration for mEGFP which were $p_{mEGFP} = 0.81 ± 0.01$ ; $B_m$ (PMT-mEGFP) = 2.32 ± 0.01; $B_d$ (PMT-mEGFP<sub>2</sub>) = 4.21 ± 0.02.

**Figure 3:**
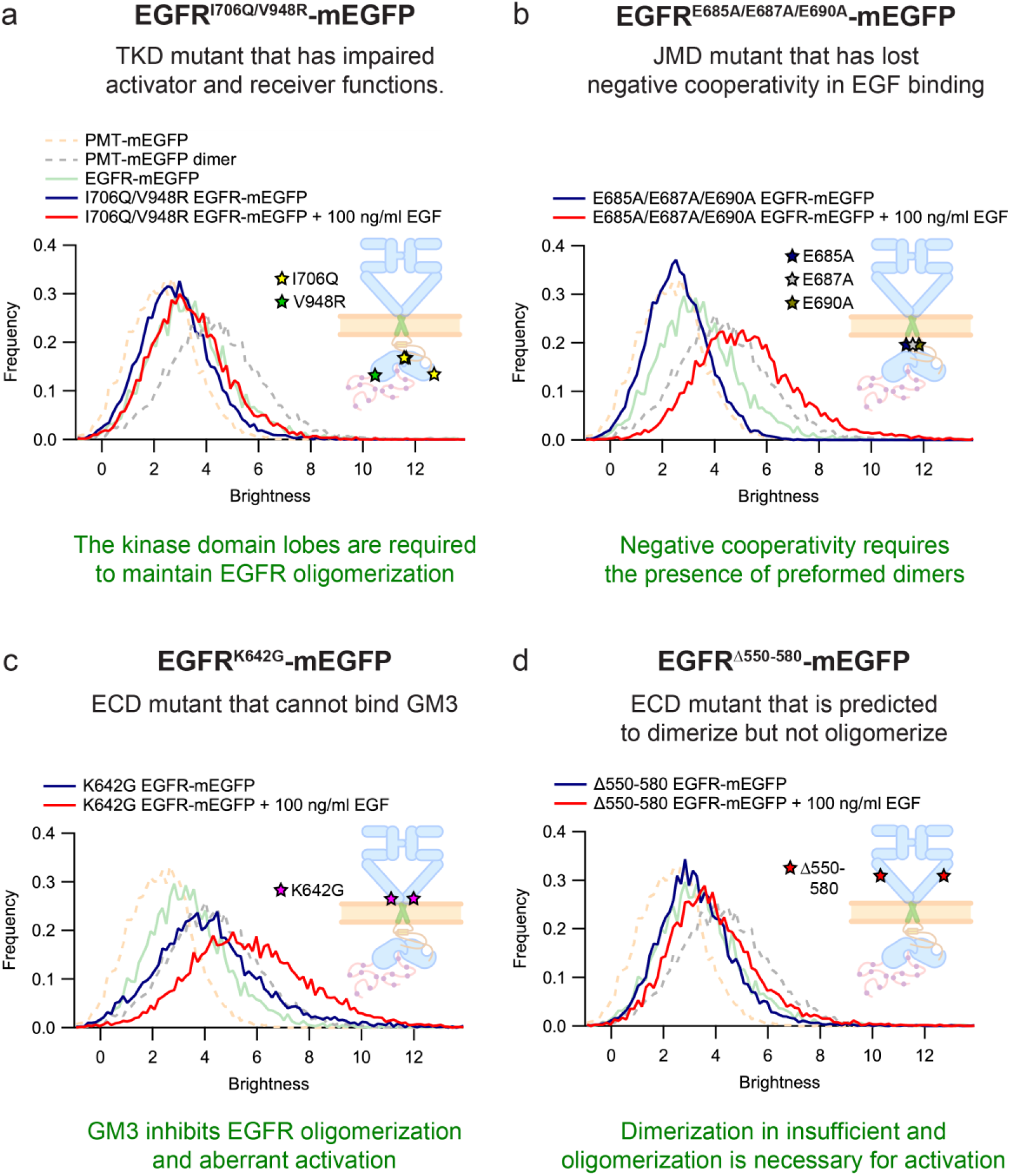
Brightness distributions of EGFR-mEGFP mutants. **(a-d)** Representative brightness histograms are shown here for the calibration controls (cream and grey dotted curves), EGFR-mEGFP in resting state (green curve), and EGFR mutants in resting state (blue curve) and after ligand stimulation (red curve). The 100 ng/ml EGF treatment curves for EGFR^E685A/E687A/E690A^-mEGFP and EGFR^K642G^-mEGFP have brightness values higher than the dimer calibration control that indicate the formation of higher-order oligomers.

#### EGFR^I706Q/V948R^-mEGFP: TKD double mutant with impaired activator and receiver function

The mutations I706Q and V948R are located on the N-terminal and C-terminal lobes, respectively, in the tyrosine kinase domain (TKD) of EGFR. These mutations disrupt the asymmetric dimer interface formed by EGFR kinase domains and cause the loss of activator and receiver function, respectively, leading to reduced phosphorylation and activation (70, 74).

The average brightness of this mutant in the resting state is 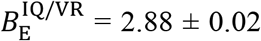 (Table 3), which is a 1.2-fold reduction as compared to *B*_E_ of EGFR-mEGFP in the resting state. The 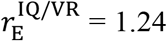 is explainable by the dimerization of 29 ± 1% of EGFR molecules. This is a 2-fold reduction as compared to the 57 ± 2% dimerization seen in the non-mutated receptor. The dimerization in the resting state is not completely abolished suggesting that other EGFR residues and/or the plasma membrane environment are also involved in formation of preformed EGFR dimers. Our results are in agreement with earlier reports using FCCS and SMP which showed that resting state oligomerization was reduced as compared to EGFR-mEGFP (26, 73).

On 10 ng/ml EGF stimulation and 100 ng/ml EGF stimulation, 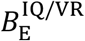 increases to 3.20 ± 0.02 and 3.36 ± 0.02 (Table 3) which are a 1.1-fold and 1.2-fold increase, respectively, from the resting state. The 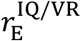 of 1.38 and 1.44 translates to 46 ± 2% and 55 ± 2% dimerization, respectively.

The phosphorylation assay showed very low phosphorylation in the resting state (Figure S8, lane 9). Unlike in the case of EGFR-mEGFP (Figure S8, lanes 5 and 6), there was also no increase in phosphorylation observed after 100 ng/ml EGF stimulation (Figure S8, lane 10) (70, 73)). The results imply that the N-terminal activator and C-terminal receiver lobes are involved in the formation of EGFR dimers in both resting and activated state.

#### EGFR^E685A/E687A/E690A^-mEGFP: JM-A triple mutant lacking negative cooperativity in EGF binding

EGFR exhibits negative cooperativity in EGF binding (71, 75, 76). The presence of negative cooperativity suggests the presence of preformed dimers as monomers cannot explain cooperativity. The juxtamembrane domain is crucial in maintaining negative cooperativity. Three glutamine residues in JM-A are predicted to be involved in interhelical salt bridges to stabilize the helical dimer. A triple-mutant (E685A/E687A/E690A) that removes these ionic interactions has been shown, using radioligand binding assays, to result in loss of negative cooperativity in EGF binding (71). This mutant was tested by N&B to directly observe if EGFR dimerization was affected.

The average *B* in the resting state is 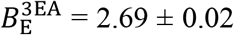 which is a 1.3-fold decrease from *B*_E_ of EGFR-mEGFP in the resting state. The 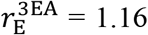 is explainable by 19 ± 1% of EGFR molecules forming dimers. This is a 3-fold reduction as compared to the 57 ± 2% dimerization seen in the non-mutated receptor. On 10 ng/ml EGF stimulation 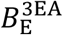 increases to 3.33 ± 0.03 (Table 3) which is a 1.2-fold increase from the resting state. The 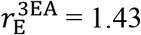 is explainable by a dimerization model and translates to 46 ± 2% EGFR molecules being present as dimers. On 100 ng/ml EGF stimulation, 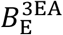 increases to 4.69 ± 0.03 (Table 3) which is a 1.7-fold increase from the resting state. The 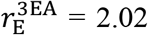 cannot be explained by a dimerization model since 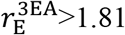. The minimum-order oligomer required is a trimer with 63 ± 2% EGFR molecules being present within trimers.

The phosphorylation assay showed no phosphorylation in the resting state (Figure S8, lane 11). On 100 ng/ml EGF stimulation increased phosphorylation was observed (Figure S8, lane 12), similar to the case of EGFR-mEGFP (Figure S8, lanes 5 and 6). The results imply that the three JM-A residues are involved in the formation of preformed dimers and mutating these residues destabilises EGFR dimers. Thus, negative cooperativity is linked to the presence of preformed dimers and the loss in negative cooperativity as observed earlier by radioligand binding assays is validated by the quantification in this study. There is ~20% dimerization still present in the resting state which indicates the involvement of other EGFR residues and/or plasma membrane components in maintaining EGFR preformed dimers. Furthermore, the EGFR molecules are able to oligomerize and be phosphorylated after EGF stimulation. This suggests these JM-A residues either do not interfere with the formation of activated EGFR dimers or that their effects are overcome by other interactions in the EGF-bound receptor.

#### EGFR^K642G^-mEGFP: ECD mutant insensitive to GM3 inhibition

The ganglioside GM3 has been shown by cross-linking, western-blot phosphorylation assays and FRET to bind to EGFR and suppress oligomerization and autophosphorylation in the absence of EGF, thereby preventing aberrant activation of the receptor (72, 77–80). The GM3-EGFR interaction is mediated through a membrane proximal lysine located in the subdomain IV of the extracellular domain (ECD). Mutation of this residue to glycine (K642G) was shown to remove GM3 binding to EGFR by simulations (79) and in phase-separated liquid-disordered (Ld)/liquid-ordered (Lo) proteoliposomes (72). To complement these previous biochemical studies on purified systems, we tested this mutant by N&B in CHO-K1 cells (GM3 makes up 0.30 ± 0.15% of total phospholipid content in this cell line (81)) to determine whether GM3-EGFR interactions are involved in EGFR clustering in live cells.

The average *B* in the resting state is 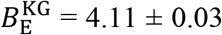 (Table 3) which is a 1.2-fold increase from *B*_E_ of EGFR-mEGFP in the resting state. The 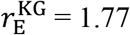 is explained by the dimerization of 95 ± 3% EGFR molecules. Upon stimulation using 10 ng/ml and 100 ng/ml EGF 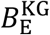 increased to 4.73 ± 0.03 and 5.53 ± 0.06 (Table 3), respectively. This represents a 1.2-fold and 1.4-fold increase from the resting state, with 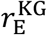 being 2.04 and 2.38, respectively. The minimum-order oligomer required to explain the data is a trimer. Using MOM, 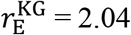 and 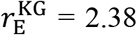 is explained by the trimerization of 64 ± 2% and 85 ± 2% of EGFR molecules, respectively.

These results in live cells are in agreement with earlier observations in phase-separated proteoliposomes (72). The K642G mutation renders EGFR insensitive to GM3 inhibition and causes an increase in resting state dimerization and phosphorylation. The results are also in line with the increase in dimerization observed after disruption of lipid domains, of which GM3 is presumably a component (82).

Given the variability in GM3 concentration across cell-lines, it would be interesting to investigate whether there is a negative correlation between EGFR dimerization proportions and GM3 concentrations in the resting state using N&B in the future.

The phosphorylation assay showed a small increase in phosphorylation in the resting state (Figure S8, lane 13) when compared to EGFR-mEGFP in the resting state (Figure S8, lane 5). Stimulation with 100 ng/ml EGF resulted in high phosphorylation (Figure S8, lane 14). This indicates that while release of GM3 inhibition allows EGFR to dimerize, EGF binding plays a role in forming higher-order oligomers that are necessary to properly activate the receptor. The requirement for higher-order oligomers to activate EGFR signalling has been suggested by other studies as well (28, 30–32, 34, 35, 58), and this theory is further tested by a multimerization-deficient mutant.

#### EGFR^Δ550-580^-mEGFP: Multimerization-deficient ECD deletion mutant

The residues from 550 to 580 in domain IV of EGFR have been predicted by simulations to be involved in multimerization but not dimerization of EGFR. SMP measurements in *Xenopus* oocytes revealed that several point mutations in this region resulted in reduced multimerization, reduced phosphorylation and similar dimerization as compared to wild-type EGFR (73). We created an EGFR mutant by deleting the entire region containing the residues 550 to 580 and probed it by N&B.

The average *B* in the resting state is 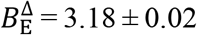 (Table 3), a 1.1-fold decrease from EGFR-mEGFP in the resting state. The 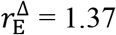 translates to 45 ± 2% of EGFR molecules forming dimers. This is a 1.3-fold reduction as compared to the 57 ± 2% dimerization seen in the non-mutated receptor. On 10 ng/ml and 100 ng/ml EGF stimulation 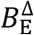 increases to 3.40 ± 0.03 and 3.64 ± 0.02 (Table 3) which are a 1.1-fold and 1.2-fold increase, respectively, from the resting state. The corresponding 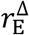 of 1.47 and 1.57 can be explained by the dimerization of 58 ± 1% and 70 ± 2% of the EGFR molecules, respectively.

The phosphorylation assay showed very low levels of phosphorylation in the resting state (Figure S8, lane 15). On 100 ng/ml EGF stimulation no increase in phosphorylation was observed (Figure S8, lane 16), unlike in the case of EGFR-mEGFP (Figure S8, lanes 5 and 6).

The average dimerization of the mutant (45 ± 2%) is less than that of EGFR-mEGFP (57 ± 2%). This reduction was also seen using SMP in an earlier study where computational modelling indicated that the formation of higher-order preformed oligomers is disrupted but dimerization is mostly unaffected (73). Moreover, the results of EGF stimulation for this mutant can be explained using an MOM model of order 2, unlike the non-mutated receptor which required order 3. This indicates that while some dimer formation probably occurred higher-order oligomerization could not be induced by EGF stimulation. Furthermore, no increase in phosphorylation is observed after 100 ng/ml EGF stimulation.

Put together the data suggests that the region containing residues 550 to 580 is involved in multimerization of EGFR and dimer formation alone is insufficient to phosphorylate EGFR. This points towards a model where EGFR multimerization is necessary to activate the receptor through phosphorylation and initiate downstream signalling. This is consistent with other studies that have also reported that dimers are not sufficient and higher-order oligomers like tetramers are necessary for EGFR signalling (28, 30–32, 34, 35, 58). It must be noted that we only report reduced oligomerization and phosphorylation in this study. The effect on actual receptor activation needs further experimental evidence through observation of recruited downstream proteins.

In summary, EGFR oligomerization is influenced by both its structural features and the plasma membrane environment, and we have quantified the oligomerization using camera-based N&B. In the resting state, EGFR molecules are present as monomers and dimers, and we estimate the dimer fraction to be comprised of ~60% of the receptor population. That said we cannot exclude the presence of a small amount of higher-order oligomers as we have shown in a recent study (27). Other previous studies (5, 26, 60–65) have shown that cholesterol- and sphingolipid-rich lipid domains trap a fraction of the EGFR population and prevent aberrant clustering and activation of the receptor in the resting state. Our quantification shows that disrupting the lipid domains through either cholesterol depletion or sphingolipids removal increases EGFR dimerization. On the other hand, cytoskeletal disruption does not affect EGFR dimerization.

Residues I706 and V948 are located in the activator and receiver lobes in the TKD, respectively, and are required for EGFR phosphorylation (70, 74) and are linked to EGFR dimerization (26, 73). The result here show that they stabilise the dimeric form of the receptor both before and after activation. Mutations to residues E685, E687 and E690 in the JM-A domain show reduced dimerization corroborating their involvement in negative cooperativity in ligand binding (71). Furthermore, as observed in proteoliposomes earlier, we show in live cells that abolishing GM3 interaction with EGFR through mutation of K642 in the ECD (72) increases receptor dimerization and phosphorylation, suggesting that GM3 acts in an inhibitory fashion to prevent aberrant receptor dimerization and phosphorylation (77–80). Finally, we also confirm the predictions from simulations that the structural requirements of dimerization differ from those of multimerization and involve different regions of the ECD. For EGFR phosphorylation (and presumably activation), dimerization is insufficient and oligomerization is necessary.

## CONCLUSION

The determination of membrane protein oligomerization is an important task in fluorescence microscopy, which is complicated by the photophysical properties of fluorophores, resulting in dark states, and the labelling process which is not complete. The measurement of oligomerization therefore requires careful calibration. Here we optimized N&B on a TIRFM setup using EMCCD cameras. We determined the probability of fluorescence of some of the most commonly used fluorescent proteins within the system with results that are consistent with confocal N&B measurements (18). Camera based N&B yields diffraction-limited oligomerization maps of ~1000 μm^2^ obtained in a measurement time of 20 s, and almost real-time analysis (~1 s) using GPU-based parallel processing, suitable for live cell investigations. Using optimized and calibrated camera-based N&B to investigate EGFR oligomerization, we determined that it is the result of a complex interplay of intrinsic structural features of the receptor and extrinsic factors including cell-membrane organization and composition. N&B has future applications in investigating clustering in live cells for other membrane proteins, and can be extended to other camera-based imaging modalities, other cameras (27), and other samples.

## AUTHOR CONTRIBUTIONS

T.W. and H.B. conceived and designed the study. T.W. supervised the study and provided the resources. H.B. and C.J.H.G. performed the experiments. H.B. and J.S. formulated the N&B equations. H.B. analyzed the data. H.B., J.S., C.G.H., and T.W. wrote the manuscript.

## ACKNOWLEDGEMENTS

T.W. gratefully acknowledges funding from the Singapore Ministry of Education (MOE2016-T2-2-121). H.B. is the recipient of a research scholarship of the National University of Singapore. The authors thank Sonia Monti for help with preparing the EGFR figures.

