## Supplementary Information for "The dependence of EGFR oligomerization on environment and structure: A camera-based N&B study"

#### for

### 1 SUPPLEMENTAL THEORY

#### 1.1 G1 analysis

Photon shot noise affects the signal variance. As shot noise is not correlated at different times the covariance between neighbouring frames was used instead as an approximation for the variance.

$$\sigma^2 = \langle (I - \langle I \rangle)^2 \rangle = \langle \delta I(t)^2 \rangle \approx \langle \delta I(t) \delta I(t + \Delta t) \rangle \tag{S1}$$

where $\delta I(t)$ is the fluorescence intensity fluctuations at time $t$, and $\delta I(t + \Delta t)$ is the fluorescence intensity fluctuations at time $t + \Delta t$ ($\Delta t$ is the exposure time of a frame). This approximation was defined as G1 analysis by Unruh et al. (1). The only condition to be satisfied to apply the G1 analysis is that the exposure time of each frame must be faster than the diffusion time of a molecule through a pixel (2).

#### 1.2 Fixed-pattern noise correction

The background correction to eliminate fixed-pattern noise is performed pixel-wise:

$$I_{ijk,corrected} = I_{ijk} - offset_{ijk} = I_{ijk} - \frac{1}{n} \sum_{k=1}^{n} I_{ijk,dark} \tag{S2}$$

where $I_{ijk,corrected}$ is the corrected pixel intensity in row $i$, column $j$ and frame $k$ of the cell image stack, $I_{ijk}$ is the original pixel intensity of that same pixel in the cell image stack, $n$ is the total number of frames in the image stack, and $I_{ijk,dark}$ is the pixel intensity of that same pixel in the dark image stack.

#### 1.3 Pooled brightness error calculation

The pooled standard error of the mean (SEM) for the brightness values (Eq. 3) is calculated as (3):

$$\Delta B_{pooled} = \left[ \frac{\sum_{j=1}^{N_c} \Delta B_j^2 + \sum_{j=1}^{N_c} \frac{\left(B_{mean_j} - B_{pooled}\right)^2}{n_j}}{N_c} \right]^{\frac{1}{2}} \tag{S3}$$

where $B_{pooled}$ and $\Delta B_{pooled}$ are the pooled mean brightness and pooled SEM, respectively, for a population of $N_c$ cells, $B_{mean_j}$ and $\Delta B_j$ is the mean brightness and SEM of all the pixels in cell $j$, and $n_j$ is the number of pixels in cell $j$.

#### 1.4 Error calculation for fluorescence probability $p$ of an FP

$$\Delta p = \frac{B_\mathrm{d}}{B_\mathrm{m}} \sqrt{\left(\frac{\Delta B_\mathrm{m}}{B_\mathrm{m}}\right)^2 + \left(\frac{\Delta B_\mathrm{d}}{B_\mathrm{d}}\right)^2} \tag{S4}$$

where $\Delta p$, $\Delta B_\mathrm{d}$ and $\Delta B_\mathrm{m}$ are the propagated SEM of $p$, $B_\mathrm{d}$ and $B_\mathrm{m}$ (Eq. 4), respectively.

#### 1.5 EGFR oligomerization error calculations

$$\Delta r_\mathrm{E} = \frac{B_\mathrm{E}}{B_\mathrm{m}} \sqrt{\left(\frac{\Delta B_\mathrm{m}}{B_\mathrm{m}}\right)^2 + \left(\frac{\Delta B_\mathrm{E}}{B_\mathrm{E}}\right)^2} \tag{S5}$$

$$\Delta e_{oligo} = \frac{1}{(n-1)p} \sqrt{\left(\frac{r_\mathrm{E} - 1}{p}\right)^2 \Delta p^2 + \Delta r_\mathrm{E}^2} \tag{S6}$$

where $\Delta r_\mathrm{E}$ and $\Delta e_{oligo}$ are the propagated SEM of $r_E$ (Eq. 5) and $e_{oligo}$ (Eq. 6), respectively.

#### 1.6 Error calculation of brightness of higher-order oligomers

$$\Delta B_n = \sqrt{[(n-1)p+1]^2 \Delta B_\mathrm{m}^2 + B_\mathrm{m}^2 (n-1)^2 \Delta p^2} \tag{S7}$$

where $B_n$ is the brightness of oligomer species of order $n$ (Eq. 7), and $\Delta B_n$ is the propagated SEM of $B_n$.

### 2 SUPPLEMENTAL MATERIALS AND METHODS

#### 2.1 Plasmids

The calibration controls were designed to localize different fluorescent proteins (FP) to the plasma membrane by fusing them with a plasma membrane targeting (PMT) sequence (4, 5). The construction of the plasmids PMT-mApple, PMT-mApple$_2$, EGFR-mApple, and EGFR-mEGFP are described in previous publications (3, 6). PMT-mEGFP was designed and purchased from VectorBuilder Inc. (Illinois, USA).

The plasmid "mCherry-7-EGFP" has been previously described by our group (7). The plasmids "pmVenus(L68V)-mTurquoise2" (Addgene plasmid #60493 (8)) and "pmScarlet_C1" (Addgene plasmid #85042 (9)) were gifts from Dorus Gadella. The plasmid "p3E-mKate2-myc no-pA" (Addgene plasmid #80812 (10)) was a gift from Philip Washbourne. The plasmid "mCherry2-N1" (Addgene plasmid #54517; a gift originally from Michael Davidson) was gifted to us by Salvatore Chiantia (11). mCherry, mTurquoise2, mScarlet, mKate2 and mCherry2 sequences were isolated from these plasmids, respectively, using the following procedure. Polymerase chain reaction (PCR) with suitably designed primers was used to amplify the FP sequences and add a SpeI recognition sequence at the 5' end and NheI and

HindIII recognition sequences at the 3' end. The amplified PCR products were digested with Spe I (SpeI-HF, #R3133S, New England Biolabs, Massachusetts, USA) and HindIII (HindIII-HF, #R3104S, New England Biolabs, Massachusetts, USA) restriction endonucleases to obtain the FP sequence insert. PMT-mEGFP was digested with SpeI and HindIII to generate the PMT backbone. The backbone and FP inserts were ligated using T4 DNA ligase (#M0202S, New England Biolabs, Massachusetts, USA) to create the PMT-FP plasmids. Thus, PMT-mCherry, PMT-mTurquoise2, PMT-mScarlet, PMT-mKate2 and PMT-mCherry2 plasmids were created.

PMT-$FP_2$ plasmids were constructed in a similar manner to PMT-$mApple_2$. Two separate digestions of PMT-FP were done using pairs of restriction endonucleases – NheI (NheI-HF, #R3131S, New England Biolabs, Massachusetts, USA) and HindIII to create the vector backbone, and SpeI and HindIII to create the FP sequence insert. The backbone and insert were ligated (possible because SpeI and NheI are isocaudomers) using T4 DNA ligase to create PMT-$FP_2$ plasmid. Thus, PMT-$mEGFP_2$, PMT-$mCherry_2$, PMT-$mTurquoise2_2$, PMT-$mScarlet_2$, PMT-$mKate2_2$, and PMT-$mCherry2_2$ plasmids were created.

With EGFR-mEGFP as the template and using suitable primers, the Q5 Site-Directed Mutagenesis Kit (#E0554S, New England Biolabs, Massachusetts, USA) was used to make the EGFR mutant constructs – $EGFR^{I706Q/V948R}$-mEGFP, $EGFR^{E685A/E687A/E690A}$-mEGFP, $EGFR^{K642G}$-mEGFP, and $EGFR^{\Delta550\text{-}580}$-mEGFP. In EGFR literature, two amino acid numbering systems are widely used. One system starts the numbering from the signal peptide and corresponds to the nascent protein. The other system excludes the signal peptide and corresponds to the mature protein. In this work, the nascent protein numbering is adopted.

#### 2.2 Cell culture and transfection

A detailed step-by-step protocol describing the preparation of live cell samples for fluorescence applications is available in Protocol Exchange (12) and also described briefly here. CHO-K1 cells (CCL-61; ATCC, Manassas, Virginia, USA) were cultured in Dulbecco's Modified Eagle Medium (DMEM/High glucose with L-glutamine, without sodium pyruvate – #SH30022.FS, HyClone, GE Healthcare Life Sciences, Utah, USA) supplemented with 1% penicillin-streptomycin (#15070063, Gibco, Thermo Fisher Scientific, Massachusetts, USA), and 10% foetal bovine serum (FBS; #10270106, Gibco, Thermo Fisher Scientific, Massachusetts, USA). They were grown in 25 $cm^2$ (#430639) or 75 $cm^2$ (#431464U) polystyrene cell culture flasks (Corning, New York, USA) and passaged regularly upon confluency. The cultures were maintained in a heated $CO_2$ incubator – 37°C and 5% (v/v) $CO_2$ environment (Forma Steri-Cycle $CO_2$ incubator, Thermo Fisher Scientific, Massachusetts, USA).

Cell cultures that were ~90% confluent were used for transfection. The spent media was removed and the culture flask was washed twice with 5 ml 1X PBS (phosphate-buffered saline; without $Ca^{2+}$ and $Mg^{2+}$). 2 ml TrypLE Express Enzyme (1X; #12604021, Gibco, Thermo Fisher Scientific, Singapore) was added and the flask was incubated at 37°C for 2-3 min to detach the cells. 5 ml culture media was added to the flask to inhibit trypsin and the cell suspension was centrifuged at 200 × g for 3 min. The supernatant was discarded and the cell pellet was resuspended in 5 ml PBS. The cells were counted using an automated cell counter (#TC20,

Bio-Rad, California, USA). The required number of cells (2-5 × $10^5$) was centrifuged (#5810, Eppendorf, Hamburg, Germany) at 200 × g for 3 min and the supernatant was discarded. The cell pellet was resuspended in R buffer (Neon Transfection Kit, Thermo Fisher Scientific, Massachusetts, USA). Suitable amounts of plasmids (100 ng for the Lifeact and PMT plasmids; 1 µg for the EGFR plasmids) were mixed with the cells. The cells were electroporated using Neon Transfection System (Thermo Fisher Scientific, Massachusetts, USA) according to the manufacturer's protocol (electroporation settings: pulse voltage = 1,000 V, pulse width = 30 ms, and pulse number = 2). The transfected cells were seeded onto culture dishes (#P35G-1.5-20-C, MatTek, Massachusetts, USA; or Nunc Lab-Tek II Chambered Coverglass, Thermo Fisher Scientific, Massachusetts, USA) containing DMEM (supplemented with 10% FBS; no antibiotic) and incubated at 37°C in 5% $CO_2$ environment for 36–48 hours before measurements.

The transfected cells were washed with Hank's Balanced Salt Solution (1X HBSS; containing $Ca^{2+}$ and $Mg^{2+}$ – #14025134, Gibco, Thermo Fisher Scientific, Massachusetts, USA) and DMEM not containing phenol red (#21063029, Gibco, Thermo Fisher Scientific, Massachusetts, USA) was added. The DMEM not containing phenol red is henceforth referred to as "imaging DMEM". EGFR-transfected cells were serum-starved in imaging DMEM for at least 4 hours to avoid aberrant activation of EGFR.

#### 2.3 Ligand stimulation and drug treatments

Working concentrations of the ligand and drugs were prepared in imaging DMEM. 10 ng/ml or 100 ng/ml of hEGF (#E9644, Sigma-Aldrich, Singapore) was added to the cells for 20 min to stimulate EGFR. Cells were treated with 3 µM LAT-A (#L5163, Sigma-Aldrich, Singapore) for 15 min to depolymerize the actin cytoskeleton. 3 mM MβCD (#C4555, Sigma-Aldrich, Singapore) treatment for 30 min depleted cholesterol from the cells. Depletion of sphingolipids was done by using 2 µM myriocin (#63150, Cayman Chemical, Michigan, USA) to treat the cells for 2 hours. All incubations were done at 37°C in 5% $CO_2$ environment.

#### 2.4 Western blotting to detect EGFR phosphorylation

2 × $10^5$ CHO-K1 cells each were transfected with 10 µg of the following plasmids: Wild-type-EGFR (WT-EGFR), EGFR-mEGFP, EGFR-mApple, $EGFR^{I706Q/V948R}$-mEGFP, $EGFR^{E685A/E687A/E690A}$-mEGFP, $EGFR^{K642G}$-mEGFP, and $EGFR^{\Delta550\text{-}580}$-mEGFP. A mock transfection without any plasmid was also done. The transfected cells in each case were divided equally into two 10 cm cell culture dishes (#353003, Corning, New York, USA) and incubated at 37°C with 5% $CO_2$ for 48 hours. The cells were then washed with 1X HBSS and serum-starved in imaging DMEM for 4 hours. Following this one dish for each sample was treated with 100 ng/ml EGF for 20 min.

A western blot kit (#12957, Western Blotting Application Solutions Kit, Cell Signaling Technology, Massachusetts, USA) was used for performing the western blots as per the manufacturer's protocol. All the cells were lysed using the provided cell lysis buffer and the cell extracts were sonicated using an ultrasonicator (VC 505, Sonics, Connecticut, USA). Sonication improves extraction of plasma membrane-bound proteins like EGFR and shears nuclear chromatin to reduce sample viscosity. The denatured proteins in the samples were separated by sodium dodecyl sulphate–polyacrylamide gel electrophoresis (SDS-PAGE). A 10–250 kDa protein ladder (#1610373, Precision Plus Protein All Blue Prestained Protein Standards, Bio-Rad, California, USA) was loaded along with the samples. Two 4-20% precast polyacrylamide gels (#4561093, Mini-PROTEAN TGX Precast Protein Gels (10-well, 30 µl), Bio-Rad, California, USA) were used in parallel – one gel was for probing with total EGFR primary antibody and the other gel with phosphorylated EGFR primary antibody.

The protein bands were wet-transferred from each PAGE gel to a nitrocellulose membrane (0.2 µm pore size; #12369, Cell Signaling Technology, Massachusetts, USA). In each sample set of two membranes, one membrane was incubated in a primary antibody solution containing total EGFR polyclonal antibody (#2232S, Cell Signaling Technology, Massachusetts, USA) and β-actin polyclonal antibody (#4967S, Cell Signaling Technology, Massachusetts, USA). The other membrane was incubated in a primary antibody solution containing phospho-EGFR (Y1173) monoclonal antibody (#4407S, Cell Signaling Technology, Massachusetts, USA) and β-actin polyclonal antibody. β-actin, a housekeeping protein, levels were used to normalize cell numbers in the samples. The secondary antibody (#7074S, Cell Signaling Technology, Massachusetts, USA) used was an IgG antibody conjugated to horseradish peroxidase (HRP).

An enhanced chemiluminescent substrate (ECL) solution (#6883, SignalFire ECL Reagent, Cell Signaling Technology, Massachusetts) was used and the chemiluminescence was detected using an imager (ImageQuant LAS 4000, GE Healthcare Bio-Sciences AB, Uppsala, Sweden) equipped with a CCD camera. The CCD camera was operated after cooling to -25°C and the images were saved as 16-bit tiff files.

#### 2.5 Calculation of EGFR phosphorylation levels

The EGFR phosphorylation levels were calculated from the western blot images as follows (6). The images were loaded in Fiji and the intensity counts were noted for the bands and the background (area on membrane with no bands).

$$\langle I_{band,corrected} \rangle = \langle I_{band} \rangle - \langle I_{background} \rangle \tag{S8}$$

where $\langle I_{band,corrected} \rangle$ is the average corrected intensity of the band, $\langle I_{band} \rangle$ is the average original intensity of the band, and $\langle I_{background} \rangle$ is the average intensity of the background (area on membrane with no bands).

To account for differences in cell numbers, the β-actin bands for corresponding lanes in the total EGFR and phosphorylated EGFR blots were normalized.

$$\langle I_{band,corrected} \rangle_{\beta,pE(nor)} = \frac{\langle I_{band,corrected} \rangle_{\beta,pE}}{\langle I_{band,corrected} \rangle_{\beta,tE}} \tag{S9}$$

where $\langle I_{band,corrected}\rangle_{\beta,pE(nor)}$ is the normalized average intensity of β-actin band in phosphorylated EGFR blot, $\langle I_{band,corrected}\rangle_{\beta,pE}$ is the original average intensity of β-actin band in phosphorylated EGFR blot, and $\langle I_{band,corrected}\rangle_{\beta,tE}$ is the original average intensity of β-actin band in total EGFR blot.

The EGFR bands for corresponding lanes in the total EGFR and phosphorylated EGFR blots were normalized to their respective normalized β-actin bands.

$$\langle I_{band,corrected}\rangle_{E,tE(nor)} = \frac{\langle I_{band,corrected}\rangle_{E,tE}}{\langle I_{band,corrected}\rangle_{\beta,tE}} \tag{S10}$$

$$\langle I_{band,corrected}\rangle_{E,pE(nor)} = \frac{\langle I_{band,corrected}\rangle_{E,pE}}{\langle I_{band,corrected}\rangle_{\beta,pE(nor)}} \tag{S11}$$

where $\langle I_{band,corrected}\rangle_{E,tE(nor)}$ is the normalized average intensity of EGFR band in total EGFR blot, $\langle I_{band,corrected}\rangle_{E,tE}$ is the original average intensity of EGFR band in total EGFR blot, $\langle I_{band,corrected}\rangle_{E,pE(nor)}$ is the normalized average intensity of EGFR band in phosphorylated EGFR blot, and $\langle I_{band,corrected}\rangle_{E,pE}$ is the original average intensity of EGFR band in phosphorylated EGFR blot.

The phosphorylation was quantified as the ratio of the average intensities of the phosphorylated EGFR band and corresponding total EGFR band.

$$Phosphorylation = \frac{\langle I_{band,corrected}\rangle_{E,pE(nor)}}{\langle I_{band,corrected}\rangle_{E,tE(nor)}} \tag{S12}$$

Note that these phosphorylation values do not report absolute phosphorylation levels. To get an estimate of relative differences we normalize and report the phosphorylation relative to the 100 ng/ml EGF stimulation of EGFR-mEGFP.

$$Relative\ phosphorylation\ (rp) = \frac{Phosphorylation_{sample}}{Phosphorylation_{EGFR-mEGFP+100ng/ml\ EGF}} \tag{S13}$$

### 3 SUPPLEMENTAL TABLES

**Table S1: EGFR dimerization in resting state as reported by different studies**

| Study | Technique | Cell line | EGFR dimerization in resting state |
|---|---|---|---|
| Moriki et al. (13) | Cross-linking | B82 | 71% |
| | | A431 | 68% |
| Kozer et al. (14) | Homo-FRET[a] | BaF/3 | 93-95% |
| Saffarian et al. (15) | FIDA[b] | CHO | >50% |
| Bader et al. (16), Hofman et al (17) | Homo-FRET[a] | NIH-3T3 | 40% |
| Nagy et al. (18) | N&B[c] | CHO | 30% |
| Kluba et al. (19) | N&B[c] | CHO-K1 | 30% |
| Zanetti-Domingues et al. (20) | FLImP[d] | CHO | 65% |
| Liu et al. (21) | FCCS[e] | CHO-K1 | 68% |
| Ma et al. (22) | FCCS[e] | CHO-K1 | 65% |
| Yavas et al. (23) | FCCS[e] | CHO-K1 | 50-70% |

[a]FRET – Förster resonance energy transfer
[b]FIDA – Fluorescence intensity distribution analysis
[c]N&B – Number and Brightness analysis
[d]FLImP – Fluorophore localization imaging with photobleaching
[e]FCCS – Fluorescence cross-correlation spectroscopy

### 4 SUPPLEMENTAL FIGURES

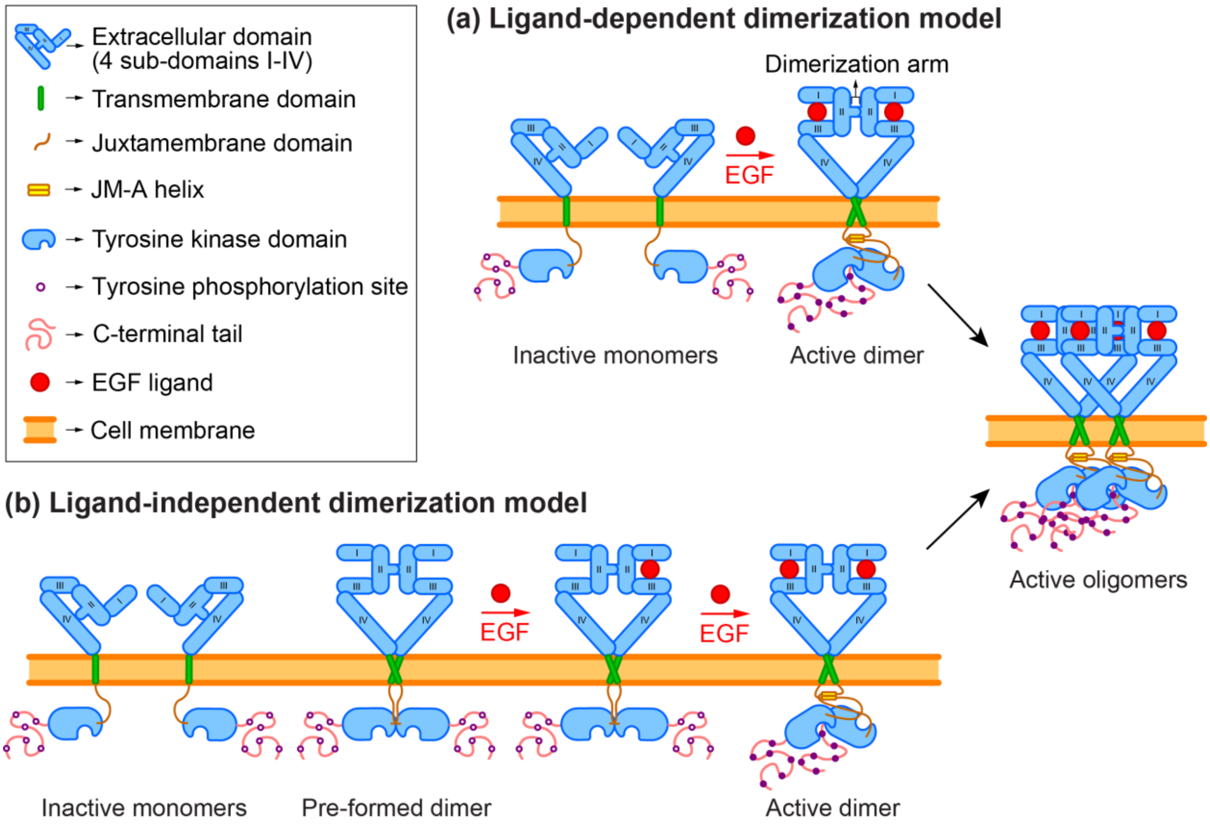


**Figure S1: EGFR activation models.** Schematic showing the ligand-dependent and ligand-independent models of EGFR activation. **(a)** In the ligand-dependent model EGFR molecules are present as inactive tethered monomers on the plasma membrane. Upon EGF binding receptors adopt an extended conformation and dimerize with other EGFR partners. The dimer partners phosphorylate each other and thereby become active to initiate downstream signalling. **(b)** In the ligand-independent dimerization model, both monomers and dimers of EGFR are present in an inactive state. Upon EGF binding receptors form active dimers for signal transduction. Some studies have suggested that dimerization is not sufficient for EGFR signalling and higher-order oligomerization (like tetramers) is required to activate the receptor.

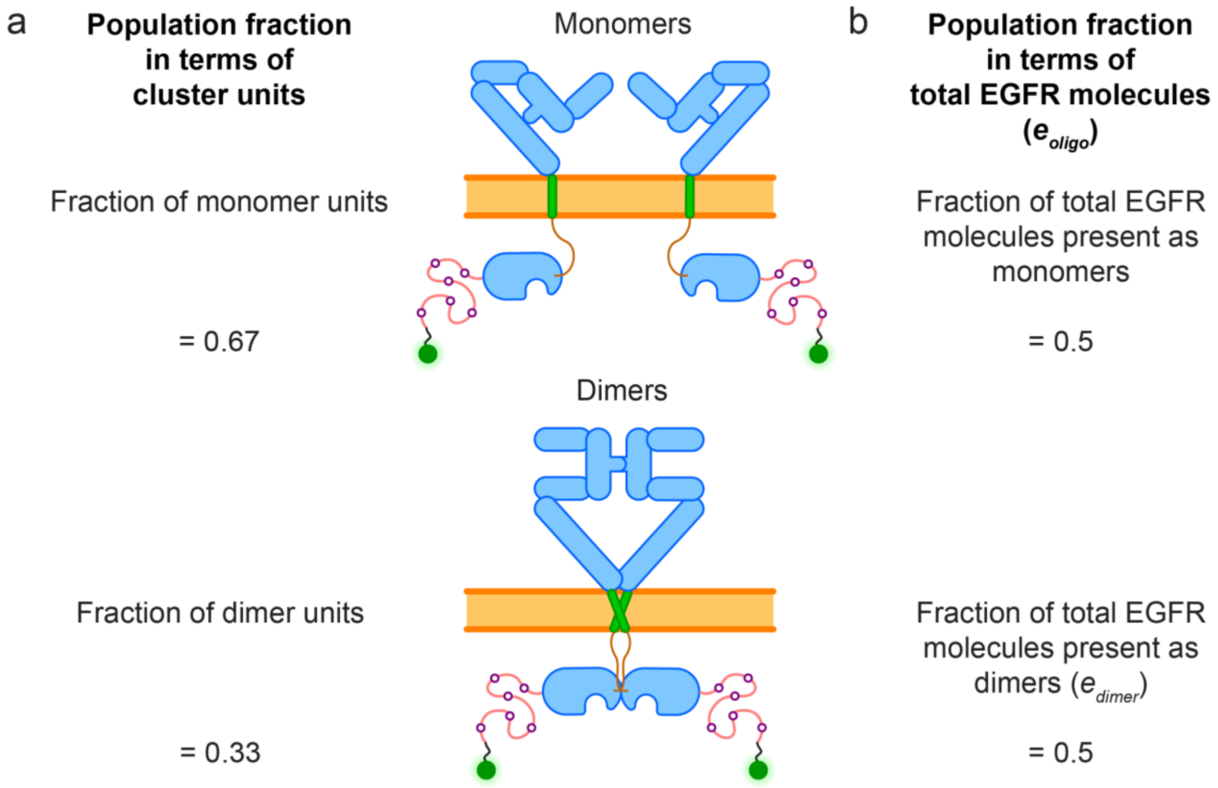


**Figure S2: Terminologies used to denote EGFR oligomer fraction.** For example, if there are four molecules of EGFR and two of them associate as dimers, there are two monomer units and one dimer unit. **(a)** If the population fraction is expressed in terms of the clustering units, the populations consists of 67% monomers and 33% dimers. **(b)** If the population fraction is expressed in terms of total EGFR molecules present in each clustering unit ($e_{oligo}$; Eq. 6) 50% of EGFR molecules are present as monomers, and another 50% are present as dimers. In literature both conventions are used and convey the same information. However, it can cause confusion if the convention used in not specified. This article follows terminology (b) to denote EGFR population fractions.

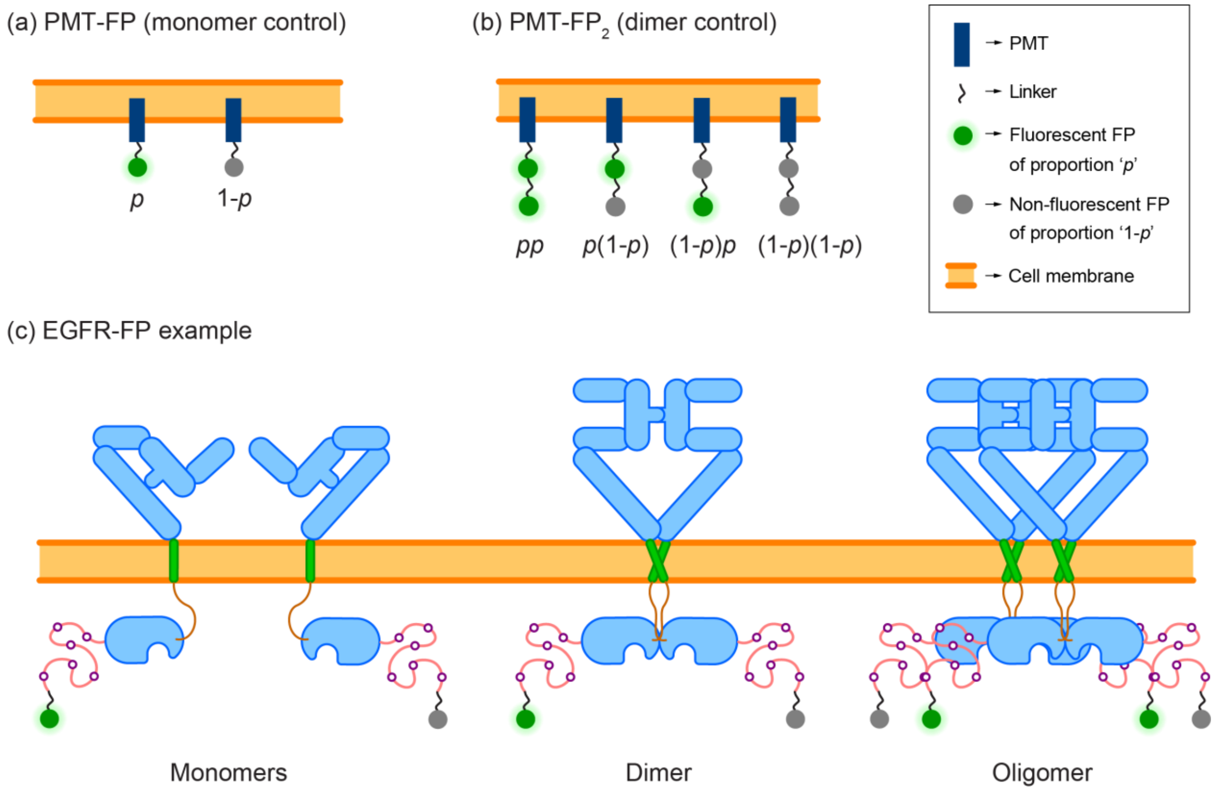


**Figure S2: Brightness calibration for N&B. (a)** A monomeric FP fused to a plasma membrane targeting (PMT) sequence serves as the monomer control. There are two population fractions in this control – fluorescent FPs of proportion *p*, and non-fluorescent FPs of proportion 1-*p*. **(b)** A tandem dimer FP fused to the PMT sequence serves as the dimer control. There are four population fractions in this control, arising from the different combinations of fluorescent and non-fluorescent moieties in the tandem dimer. The proportions of the four fractions are $p^2$, $p(1-p)$, $(1-p)p$, and $(1-p)^2$. By performing a brightness calibration using the monomer and dimer controls, *p* can be estimated using Eq. 4. **(c)** The EGFR-FP population consists of monomers, dimers, and oligomers. The quantification of these fractions requires *p* to be incorporated into the calculations (Eq. 6).

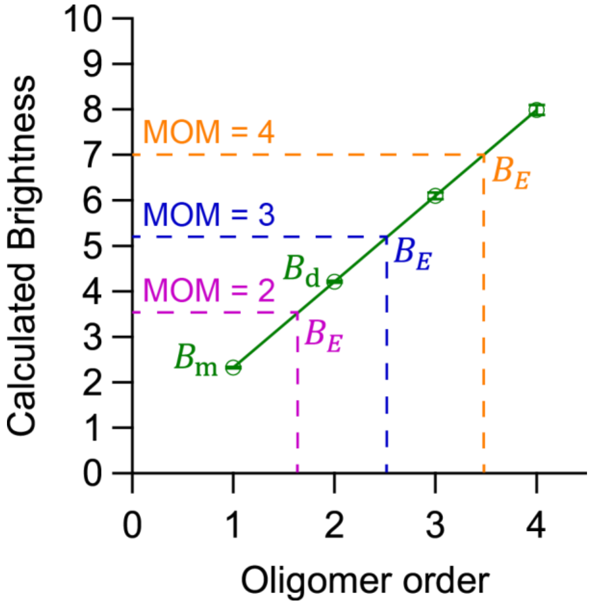


**Figure S4: Brightness scaling of oligomers.** The scaling of expected brightness with oligomer order (green line) is shown here. The *B* values of PMT-mEGFP and PMT-$mEGFP_2$ are experimentally determined (N = 3 cells). The *B* values of trimer and tetramer are estimated theoretically using Eq. 7. The minimum-order oligomer ($n_{mom}$) required can be calculated using Eq. 8. The magenta, blue and orange dotted lines show example $B_E$ values that according to the MOM require at minimum the presence of dimers, trimers and tetramers, respectively, to explain the data. The data points with error bars represent mean ± SEM.

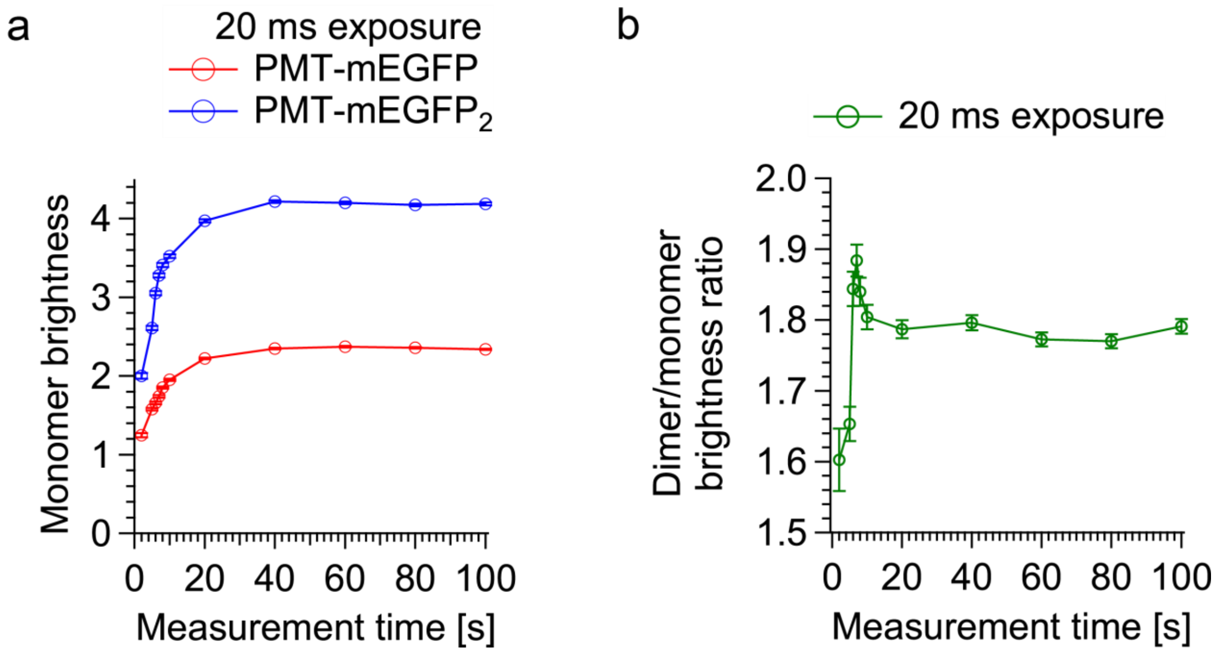


**Figure S5: Optimization of acquisition parameters for N&B.** The effect of varying the total measurement time (from 2 to 100 s) on **(a)** the brightness of the calibration controls and **(b)** the brightness ratio of PMT-mEGFP$_2$/PMT-mEGFP is plotted here. 20 ms exposure time is used. The ratio increases initially and peaks around 6 s of measurement time. It then drops and stabilises at ~1.8 from 10 s onwards. The initial increase is due to the fact that since the dimer control has two FP moieties it is brighter and achieves sufficient SNR with less measurement time as compared to the monomer control. At those times the monomer control might not have sufficient SNR and its brightness is underestimated, thus overestimating the brightness ratio. Once the monomer control also has achieved sufficient SNR the ratio stabilises (from 10 s onwards). Therefore, we use 20 s measurement time for stable brightness ratios in this work. Each point is an average of 3 different cell measurements for both PMT-mEGFP and PMT-mEGFP$_2$. The number of pixels in each cell is different due to differential intensity filtering in each case, but each cell had at least 4,000 and at most 9,000 valid pixels. The data points with error bars represent mean ± SEM.

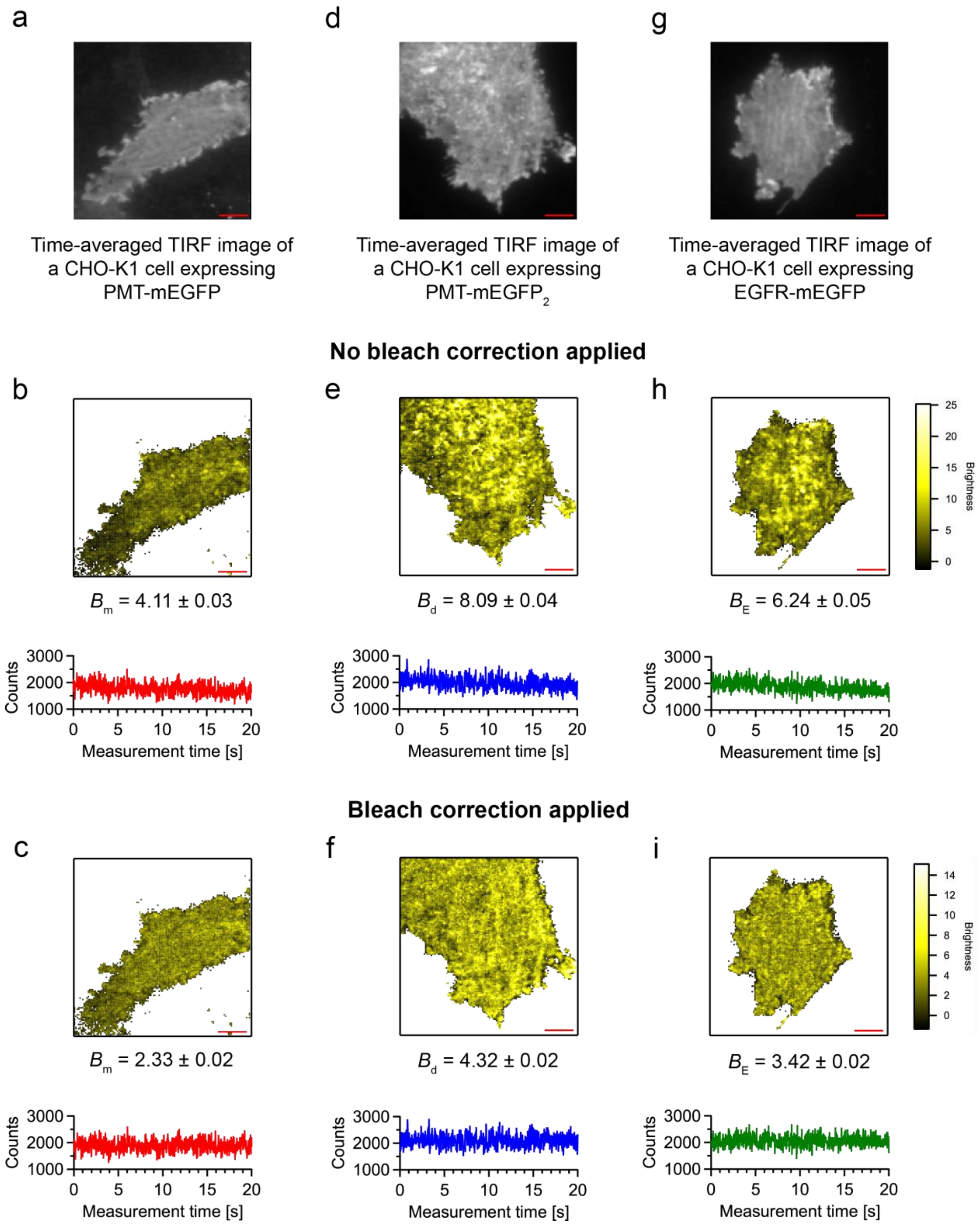


**Figure S6: Effect of photobleaching on N&B. (a)** Time-averaged TIRF image of a CHO-K1 cell expressing PMT-mEGFP. **(b)** *B* map of the cell in image (a) generated without applying bleach correction. The average brightness of the cell $B_m$ = 4.11 ± 0.03. The bottom panel shows an example intensity trace (red) from this cell depicting the decrease in intensity with time due to photobleaching. **(c)** *B* map of the cell in image (a) generated after applying bleach correction. The average brightness of the cell $B_m$ = 2.33 ± 0.02. The bottom panel shows the same intensity trace (red) in image (b) after bleach correction. The average intensity is now stationary across the measurement time. **(d)** The time-averaged TIRF image of a CHO-K1 cell expressing PMT-mEGFP$_2$. **(e)** *B* map of the cell in image (d) generated without applying bleach correction. The average brightness of the cell $B_d$ = 8.09 ± 0.04. The bottom panel shows an example intensity trace (blue) from this cell. **(f)** *B* map of the cell in image (b)

generated after applying bleach correction. The average brightness of the cell $B_d$ = 4.32 ± 0.02. The bottom panel shows the same intensity trace (blue) in image (d) after bleach correction. **(g)** Time-averaged TIRF image of a CHO-K1 cell expressing EGFR-mEGFP. **(h)** *B* map of the cell in image (g) generated without applying bleach correction. The average brightness of the cell $B_d$ = 6.24 ± 0.05. The bottom panel shows an example intensity trace (green) from this cell. **(i)** *B* map of the cell in image (b) generated after applying bleach correction. The average brightness of the cell $B_d$ = 3.42 ± 0.02. The bottom panel shows the same intensity trace (green) in image (g) after bleach correction. This figure illustrates how photobleaching is undesirable since it breaks the assumption of stationary mean and overestimates the variance. This results in overestimation of *B* values and underestimation of *N*. All the B values given here are mean ± SEM. The scale bars in red represent 5 µm.

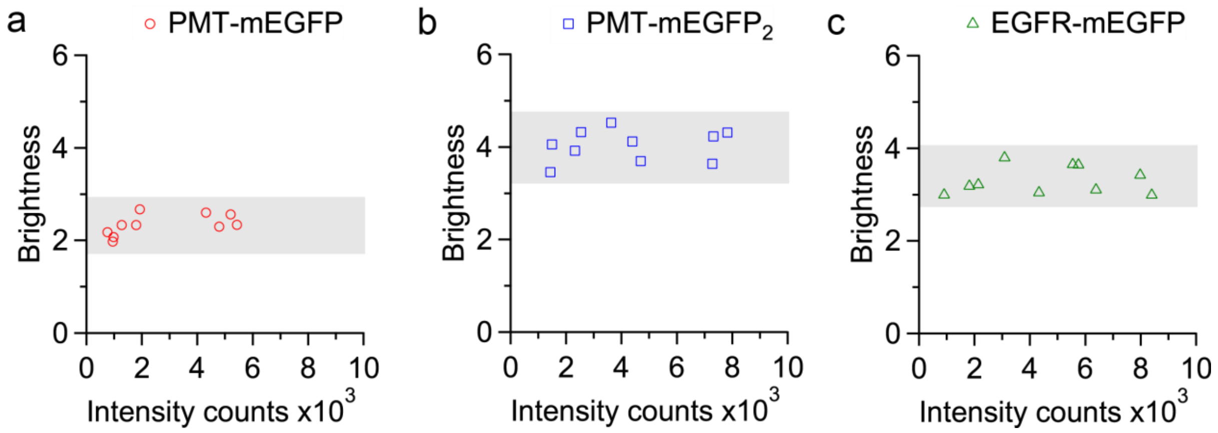


**Figure S7: *B* values show no dependence on protein expression levels.** The *B* values for 10 cells each are plotted against the intensity counts for **(a)** PMT-mEGFP, **(b)** PMT-mEGFP$_2$, and **(c)** EGFR-mEGFP. There is no expression dependence observed for the *B* values over an intensity range of 500 to 8000 counts for all the samples. The *B* values of EGFR-mEGFP are in between those of PMT-mEGFP and PMT-mEGFP$_2$.

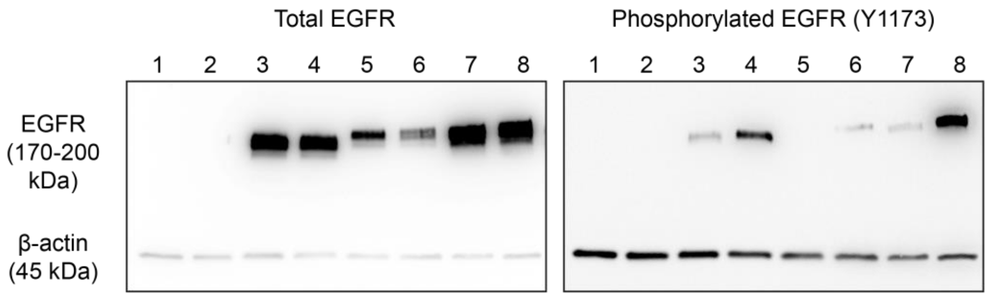


1 - Mock transfection
Relative phosphoryation = 0

2 - Mock transfection + 100 ng/ml EGF
Relative phosphoryation = 0

3 - WT-EGFR
Relative phosphoryation = 0.2

4 - WT-EGFR + 100 ng/ml EGF
Relative phosphoryation = 1.2

5 - EGFR-mEGFP
Relative phosphoryation = 0

6 - EGFR-mEGFP + 100 ng/ml EGF
Relative phosphoryation = 1

7 - EGFR-mApple
Relative phosphoryation = 0.2

8 - EGFR-mApple + 100 ng/ml EGF
Relative phosphoryation = 5

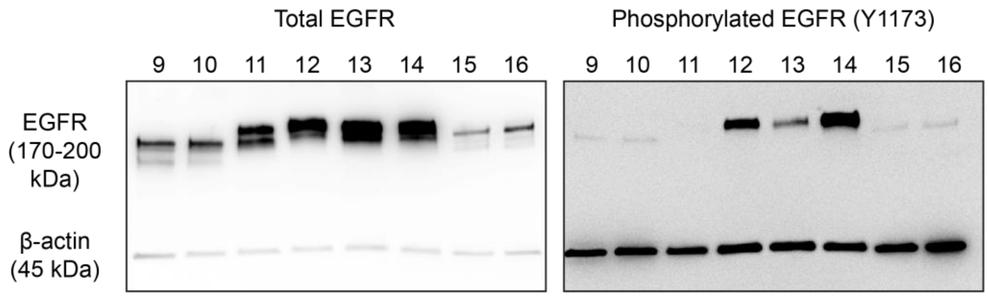


9 - $EGFR^{I706Q/V948R}$-mEGFP
Relative phosphoryation = 0.1

10 - $EGFR^{I706Q/V948R}$-mEGFP
+ 100 ng/ml EGF
Relative phosphoryation = 0.1

11 - $EGFR^{E685A/E687A/E690A}$-mEGFP
Relative phosphoryation = 0

12 - $EGFR^{E685A/E687A/E690A}$-mEGFP
+ 100 ng/ml EGF
Relative phosphoryation = 1.8

13 - $EGFR^{K642G}$-mEGFP
Relative phosphoryation = 0.4

14 - $EGFR^{K642G}$-mEGFP
+ 100 ng/ml EGF
Relative phosphoryation = 6.4

15 - $EGFR^{\Delta 550\text{-}580}$-mEGFP
Relative phosphoryation = 0.2

16 - $EGFR^{\Delta 550\text{-}580}$-mEGFP
+ 100 ng/ml EGF
Relative phosphoryation = 0.1

**Figure S3: Western blot detection of EGFR phosphorylation.** A number of samples were probed for EGFR phosphorylation. Three primary antibodies were used – a polyclonal antibody to detect total EGFR levels, a monoclonal antibody to detect phosphorylated tyrosine residue 1173 (Y1173), and a polyclonal β-actin antibody to normalize variations in cell numbers. The EGFR band is detected at ~170-200 kDa (the unlabelled receptor is typically seen at ~170 kDa; addition of an FP tag increases its molecular weight by ~27 kDa). The β-actin band is detected at ~45 kDa. Further details on the western blotting and calculation of relative phosphorylation (rp) levels are provided in Supplemental materials and methods. Each corresponding set of blots were processed in parallel. The contrast of the images has been adjusted to show all the bands, including the faint ones, clearly. Among the 16 lanes, odd-numbered lanes represent resting cells and even-numbered lanes represent cells stimulated with 100 ng/ml EGF. Multiple bands are seen in the total EGFR lane in some cases and indicates either sample degradation or post-translational modifications. The western blot was performed twice with similar results and one representative set of blots are shown here.

**(1) Mock transfection:** CHO-K1 cells transfected without using any EGFR plasmid. As expected, no EGFR presence is detected and no phosphorylation (rp = 0) is seen.
**(2) Mock transfection + 100 ng/ml EGF:** Mock transfected CHO-K1 cells that were subjected to 100 ng/ml EGF stimulation. Since no EGFR was transfected, the treatment had no effect. No EGFR is detected and no phosphorylation (rp = 0) is observed.
**(3) WT-EGFR:** CHO-K1 cells transfected with WT-EGFR not containing any fluorescent tag. The presence of EGFR is detected and some baseline phosphorylation (rp = 0.2) is seen. This sample acts as a control to verify that fluorescent labelling of EGFR does not affect its activity.
**(4) WT-EGFR + 100 ng/ml EGF:** CHO-K1 cells transfected with untagged WT-EGFR and stimulated using 100 ng/ml EGF. The presence of EGFR is detected and phosphorylation increased (1.2) as expected when compared to the resting state (lane 3).
**(5) EGFR-mEGFP:** CHO-K1 cells transfected with EGFR-mEGFP plasmid. The presence of EGFR is detected. No baseline phosphorylation (rp = 0) is seen – it is possible the lower amounts of total EGFR present as compared to lane 3 results in the baseline phosphorylation being too low and below the detection limit of the system.
**(6) EGFR-mEGFP + 100 ng/ml EGF:** CHO-K1 cells transfected with EGFR-mEGFP plasmid and stimulated with 100 ng/ml EGF. The presence of EGFR is detected and increased phosphorylation (rp = 1) is seen as compared to the resting state (lane 5). This indicates that the mEGFP tag doesn't affect the stimulation of EGFR.
**(7) EGFR-mApple:** CHO-K1 cells transfected with EGFR-mApple plasmid. The presence of EGFR is detected and some baseline phosphorylation (rp = 0.2) is seen.
**(8) EGFR-mApple + 100 ng/ml EGF:** CHO-K1 cells transfected with EGFR-mApple plasmid and stimulated with 100 ng/ml EGF. The presence of EGFR is detected and increased phosphorylation (rp = 5) is seen as compared to the resting state (lane 7). This indicates that the mApple tag used in our previous publication (3) doesn't affect the stimulation of EGFR.
**(9) EGFR$^{I706Q/V948R}$-mEGFP:** CHO-K1 cells transfected with EGFR$^{I706Q/V948R}$-mEGFP plasmid. The presence of EGFR is detected and low baseline phosphorylation (rp = 0.1) is observed.
**(10) EGFR$^{I706Q/V948R}$-mEGFP + 100 ng/ml EGF:** CHO-K1 cells transfected with EGFR$^{I706Q/V948R}$-mEGFP plasmid and stimulated with 100 ng/ml EGF. The presence of EGFR is detected and no increase in phosphorylation (rp = 0.1) is seen as compared to the resting state (lane 9).
**(11) EGFR$^{E685A/E687A/E690A}$-mEGFP:** CHO-K1 cells transfected with EGFR$^{E685A/E687A/E690A}$-mEGFP plasmid. The presence of EGFR is detected and no baseline phosphorylation (rp = 0; possibly below detection limit) is observed.
**(12) EGFR$^{E685A/E687A/E690A}$-mEGFP + 100 ng/ml EGF:** CHO-K1 cells transfected with EGFR$^{E685A/E687A/E690A}$-mEGFP plasmid and stimulated with 100 ng/ml EGF. The presence of EGFR is detected and an increase in phosphorylation (rp = 1.8) is observed as compared to the resting state (lane 11).
**(13) EGFR$^{K642G}$-mEGFP:** CHO-K1 cells transfected with EGFR$^{K642G}$-mEGFP plasmid. The presence of EGFR is detected, and the baseline phosphorylation (rp = 0.4) observed is slightly higher than the WT receptor in the resting state.
**(14) EGFR$^{K642G}$-mEGFP + 100 ng/ml EGF:** CHO-K1 cells transfected with EGFR$^{K642G}$-mEGFP plasmid and stimulated with 100 ng/ml EGF. The presence of EGFR is detected and an increase in phosphorylation (rp = 6.4) is observed as compared to the resting state (lane 13).
**(15) EGFR$^{\Delta550\text{-}580}$-mEGFP:** CHO-K1 cells transfected with EGFR$^{\Delta550\text{-}580}$-mEGFP plasmid. The presence of EGFR is detected and low baseline phosphorylation (rp = 0.2) is observed.
**(16) EGFR$^{\Delta550\text{-}580}$-mEGFP + 100 ng/ml EGF:** CHO-K1 cells transfected with EGFR$^{\Delta550\text{-}580}$-mEGFP plasmid and stimulated with 100 ng/ml EGF. The presence of EGFR is detected and no increase in phosphorylation (rp = 0.1) is observed as compared to the resting state (lane 15).

14. Kozer, N., M.P. Kelly, S. Orchard, A.W. Burgess, A.M. Scott, and A.H.A. Clayton.

2011. Differential and synergistic effects of epidermal growth factor receptor antibodies on unliganded ErbB dimers and oligomers. *Biochemistry*. 50:3581–3590.

20. Zanetti-Domingues, L.C., D. Korovesis, S.R. Needham, C.J. Tynan, S. Sagawa, S.K. Roberts, A. Kuzmanic, E. Ortiz-Zapater, P. Jain, R.C. Roovers, A. Lajevardipour, P.M.P. van Bergen en Henegouwen, G. Santis, A.H.A. Clayton, D.T. Clarke, F.L. Gervasio, Y. Shan, D.E. Shaw, D.J. Rolfe, P.J. Parker, and M.L. Martin-Fernandez. 2018. The architecture of EGFR's basal complexes reveals autoinhibition mechanisms in dimers and oligomers. *Nat. Commun.* 9.

21. Liu, P., T. Sudhaharan, R.M.L. Koh, L.C. Hwang, S. Ahmed, I.N. Maruyama, and T. Wohland. 2007. Investigation of the dimerization of proteins from the epidermal growth factor receptor family by single wavelength fluorescence cross-correlation spectroscopy. *Biophys. J.* 93:684–698.

22. Ma, X., S. Ahmed, and T. Wohland. 2011. EGFR activation monitored by SW-FCCS in live cells. *Front. Biosci. - Elit.* 3 E:22–32.

23. Yavas, S., R. Macháň, and T. Wohland. 2016. The Epidermal Growth Factor Receptor Forms Location-Dependent Complexes in Resting Cells. *Biophys. J.* 111:2241–2254.
